# Cellular and Molecular Probing of Intact Transparent Human Organs

**DOI:** 10.1101/643908

**Authors:** Shan Zhao, Mihail Ivilinov Todorov, Ruiyao Cai, Hanno Steinke, Elisabeth Kemter, Eckhard Wolf, Jan Lipfert, Ingo Bechmann, Ali Ertürk

## Abstract

Optical tissue transparency permits cellular and molecular investigation of complex tissues in 3D, a fundamental need in biomedical sciences. Adult human organs are particularly challenging for this approach, owing to the accumulation of dense and sturdy molecules in decades-aged human tissues. Here, we introduce SHANEL method utilizing a new tissue permeabilization approach to clear and label stiff human organs. We used SHANEL to generate the first intact transparent adult human brain and kidney, and perform 3D histology using antibodies and dyes in centimeters depth. Thereby, we revealed structural details of sclera, iris and suspensory ligament in the human eye, and the vessels and glomeruli in the human kidney. We also applied SHANEL on transgenic pig organs to map complex structures of EGFP expressing beta cells in >10 cm size pancreas. Overall, SHANEL is a robust and unbiased technology to chart the cellular and molecular architecture of intact large mammalian organs.

**Graphical Abstract:** 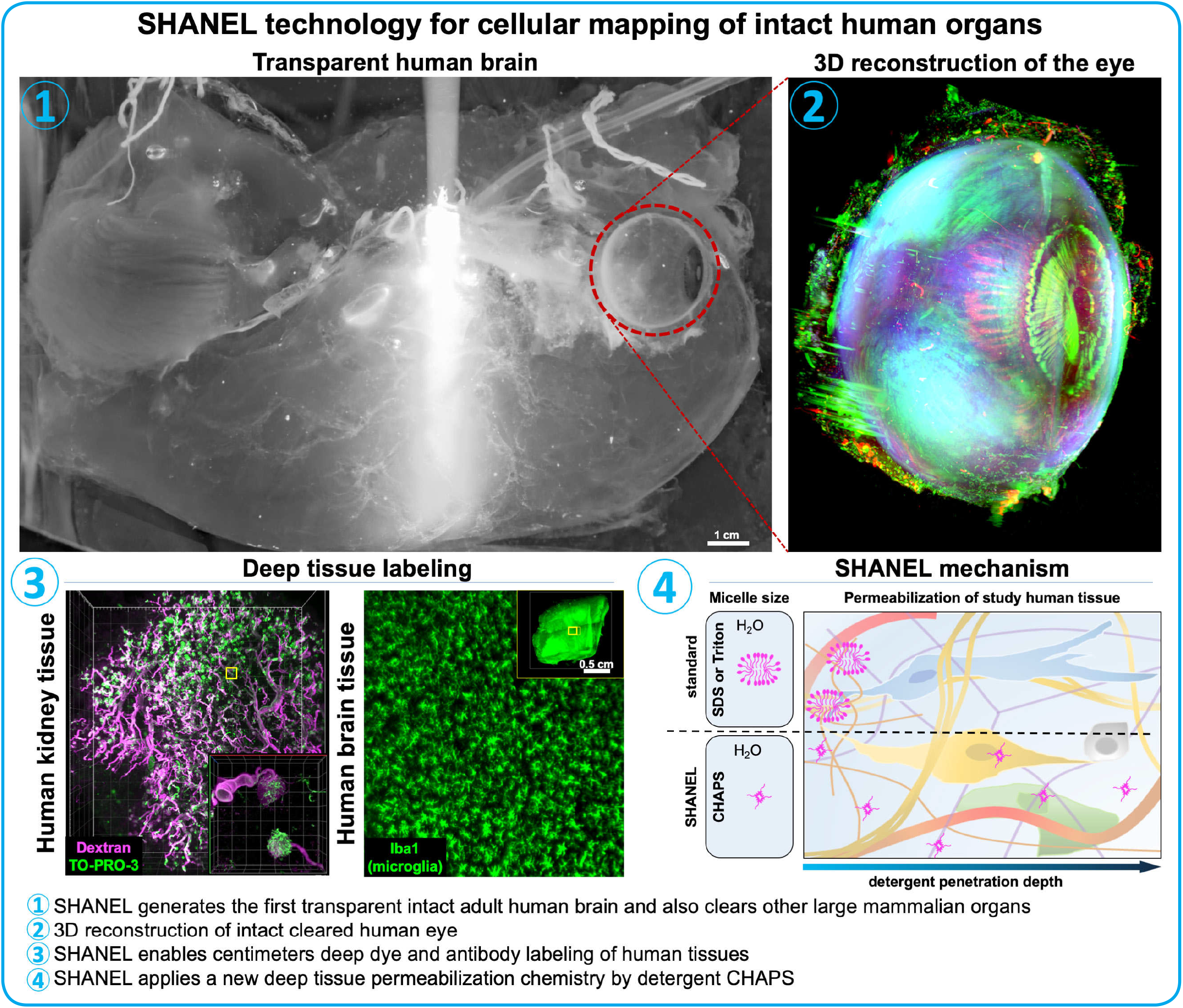

Supplementary Movies of SHANEL are available at http://discotechnologies.org/SHANEL/

## INTRODUCTION

Structural and functional mapping of human organs, especially the human brain, has been one of the flagship projects of biomedical science in the last years. Towards this goal, the US, the EU and many other countries initiated their own “human brain mapping” projects [1]. Yet, the progress has been limited especially in deciphering the anatomical complexity of human brain, mainly due to lack of scalable technologies to image the content of large human organs at the cellular level. While magnetic resonance imaging (MRI) can provide longitudinal imaging for human organs including brain and kidney, it lacks cellular resolution [2–4]. Therefore, tissue histology has been the major approach to study the molecular and cellular complexity of the human brain. Although routine standard histology is limited to tiny pieces of the human brain (typically a tissue section is ~1/10,000,000 of the whole brain volume), there still have been efforts to perform it for whole human brain mapping [5, 6]. However, slicing and imaging thousands of thin sections from a whole human brain alone may require years of labor, and subsequent reconstruction of the whole brain in 3D could be very complicated or impossible due to the numerous tissue distortions introduced by mechanical sectioning. Thus, a scalable and routine technology to enable cellular and molecular interrogation of centimeters sized human organs could substantially reduce sectioning artifacts and also overcome complications in registering large scale imaging data in 3D. Such an approach would eliminate the extremely tedious and laborious work with thousands of thin sections that is needed to map human organs.

In the last decade, emerging optical tissue clearing methods have enabled fast 3D histology on transparent specimens avoiding major pitfalls of standard histology, especially tissue sectioning [7, 8]. Furthermore, new deep tissue labeling methods were developed in combination with clearing methods to better phenotype whole rodent organs and human embryos [9–15]. Progress in optical tissue clearing firstly allowed clearing of increasingly larger rodent samples (up to whole adult rodent bodies) [16–19], then adaptation of light-sheet microscopy systems to manage imaging of whole transparent rodent bodies [14, 18, 20]. However, clearing of human organs so far has been notoriously challenging, in particular for adult human brain tissue. Recent efforts with chemical screening of thousands of compounds [21] and application of electrical field forces [22] could achieve clearing of only small pieces of human organs. For example, it took 10 months to clear a 8 mm-thick human brain specimen [23] and 3.5 months to clear a 5 mm-thick human striatum sample [24]. Furthermore, deep-tissue antibody labeling methods developed on rodent tissues also encounter hurdles to label adult human tissue thicker than 1 mm [25]. We reasoned that highly myelinated content, lipidome complexity [26], and age-related accumulation of opaque and dense molecules such as lipofuscin and non-soluble collagen [27, 28] impede penetration of chemicals deep into human organs, thereby blocking both clearing and labeling of centimeters sized specimens.

Here, we introduce SHANEL (Small-micelle-mediated Human orgAN Efficient clearing and Labeling), a new method that is driven by detergent permeabilization chemistry allowing penetration of labeling and clearing agents into centimeters-thick mammalian organs (**Fig. 1**). Our approach enables histology of human samples ranging from 1.5 cm thickness to whole adult human organs. We also show that the technology works on other large mammalian organs such as pig brain and pancreas, which can readily be labeled transgenically. Thus, the SHANEL histology pipeline presented here will pave the way for cellular and molecular mapping of whole adult human organs, including the human brain.

**Fig. 1.**
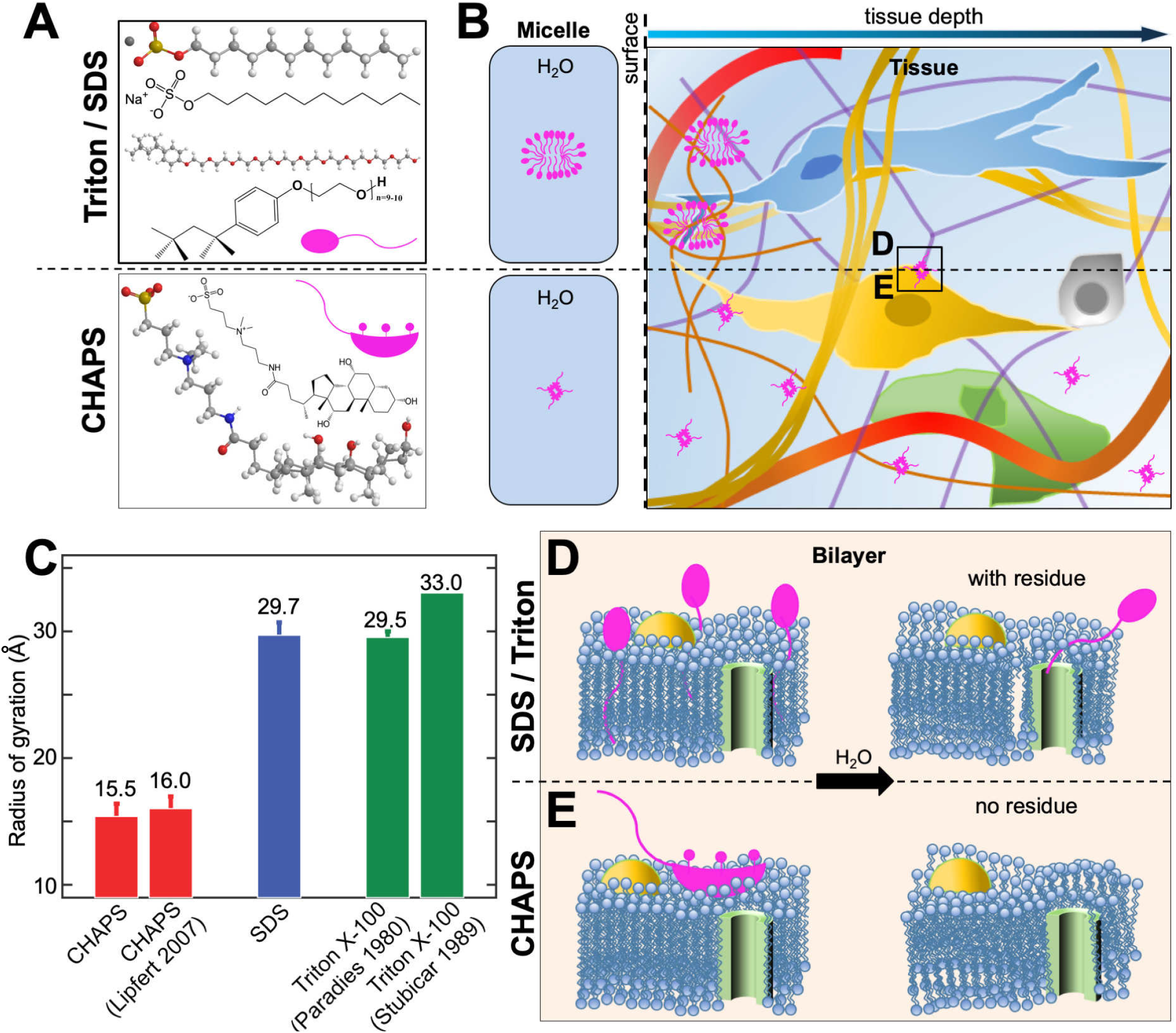
Deep tissue permeabilization by detergent CHAPS forming small micelle. **(A)** 3D molecular structural features of SDS and Triton X-100 exhibiting typical head-to-tail amphiphilicity, whereas, CHAPS exhibiting peculiar facial amphiphilicity. **(B)** Schematic diagram showing the formation of smaller micelle by facial CHAPS compared to standard detergents for tissue clearing and histology (SDS and Triton X-100). CHAPS could more efficiently and deeply permeabilize the tissue owing to its smaller micelle size. **(C)** Radii of gyration determined for CHAPS and SDS and literature values (full references are in TableS1) for *R_g_* (the size of micelle is characterized by the squared radius of gyration of the micellar core, *R_g_*^2^). Values for CHAPS and SDS represent mean and standard error from at least three independent repeats. **(D,E)** Detectable fragments of SDS and Triton X-100 are evident after washing while no residual of CHAPS is left inside of the tissue. Facial CHAPS disrupts the lipid bilayer by binding onto the surface rather than by embedding into the membrane as head-to-tail detergents, therefore CHAPS could be completely washed out.

## RESULTS

### Development of new detergent permeabilization chemistry

We hypothesized that both labeling and clearing of large and sturdy human organs require a new permeabilization chemistry allowing deep tissue penetration of molecules. Ionic SDS (Sodium dodecyl sulfate) and nonionic Triton X-100 (4-(1,1,3,3-Tetramethylbutyl)phenyl-polyethylene glycol) are commonly used detergents for tissue clearing, and they are characterized as containing typical ‘head-to-tail’ amphipathic regions. Their structural features lead to the formation of relatively large micelles suggesting that they can get stuck at the tissue surfaces, and therefore exhibit limited tissue permeabilization capacity (**Fig. 1A-C; Supporting Fig. 1; Supporting Table 1**). We anticipated that detergents forming smaller micelles would be better candidates for deep tissue permeabilization as they could penetrate more rapidly and deeply into the tissue. We identified the zwitterionic detergent 3-[(3-cholamidopropyl)dimethylammonio]-1-propanesulfonate (CHAPS), bearing a rigid steroidal structure with a hydrophobic convex side, a hydrophilic concave side (bearing three hydroxyl groups) and a sulfobetaine-type polar group. CHAPS possesses atypical ‘facial’ amphiphilicity [29]. With this peculiar structure of hydrophobic and hydrophilic faces, it has higher critical micelle concentration (CMC), smaller aggregation number, and forms much smaller micelles compared to SDS and Triton X, which could enable its rapid penetration deep into large and sturdy human organs such as the brain with densely packed extracellular matrix molecules (**Fig. 1A-C; Supporting Fig. 1; Supporting Table 1**). Furthermore, once the micelles travel into the tissue and interact with the bilayer of lipid, the facial hydrophobic side of CHAPS reclines on the bilayer surface with a larger area rather than being embedded into the lipid as head-to-tail detergents. These different interaction behaviors of the detergents suggest that detectable fragments of SDS and Triton X-100 exist after washing while no residual of CHAPS would be left inside of the tissue [30–32] (**Fig. 1D,E**). Thus, CHAPS could function as an efficient tissue permeabilization reagent by traveling throughout sturdy tissue and disrupting the dense ultrastructure, without leaving behind residual detergent fragments after wash out.

Human organs carry residual blood clots due to several hours to days of delay until organ harvest after death. Heme in the blood causes strong autofluorescence at visible wavelengths (400-700 nm) and reduces the intensity of travelling light within the tissue, thereby impeding the full transparency of cleared organs [17]. PFA-fixed blood was washed by detergent solutions, resulting in colorless supernatant and red pellet, indicating detergent alone was unable to remove the heme (**Fig. 2A**). To overcome this issue, we firstly screened CHAPS compatible chemicals to elute the heme. In particular, we focused on effective, colorless and cheap chemicals for scalability to large human organs (**Fig. 2B-E**). Our screen showed that N-methyldiethanolamine (NMDEA) was an efficient candidate when combined with CHAPS, resulting in a completely colorless pellet from PFA-fixed blood (chemical 7 in **Supporting Tables 2-3**). In addition, NMDEA was the cheapest among the screened chemicals, reducing the cost when used in large amounts for intact human organs (**Supporting Table 3**). Compared to Triton X-100 and SDS that were used in prior clearing methods as detergents, CHAPS was faster and more successful in decolorization of mouse kidney, liver, heart, spleen and brain, suggesting a better performance for large human organs (**Fig. 2F**). Moreover, protein loss assay indicated the superior retention of endogenous biomolecules with CHAPS, assuring a more reliable molecular investigation of intact organs (**Fig. 2G**). Thus, we anticipated that CHAPS, by forming small micelles, could completely diffuse through intact large mammalian organs and ameliorate tissue meshes, leaving behind a fully permeabilized biological tissue for cellular and molecular phenotyping.

**Fig. 2.**
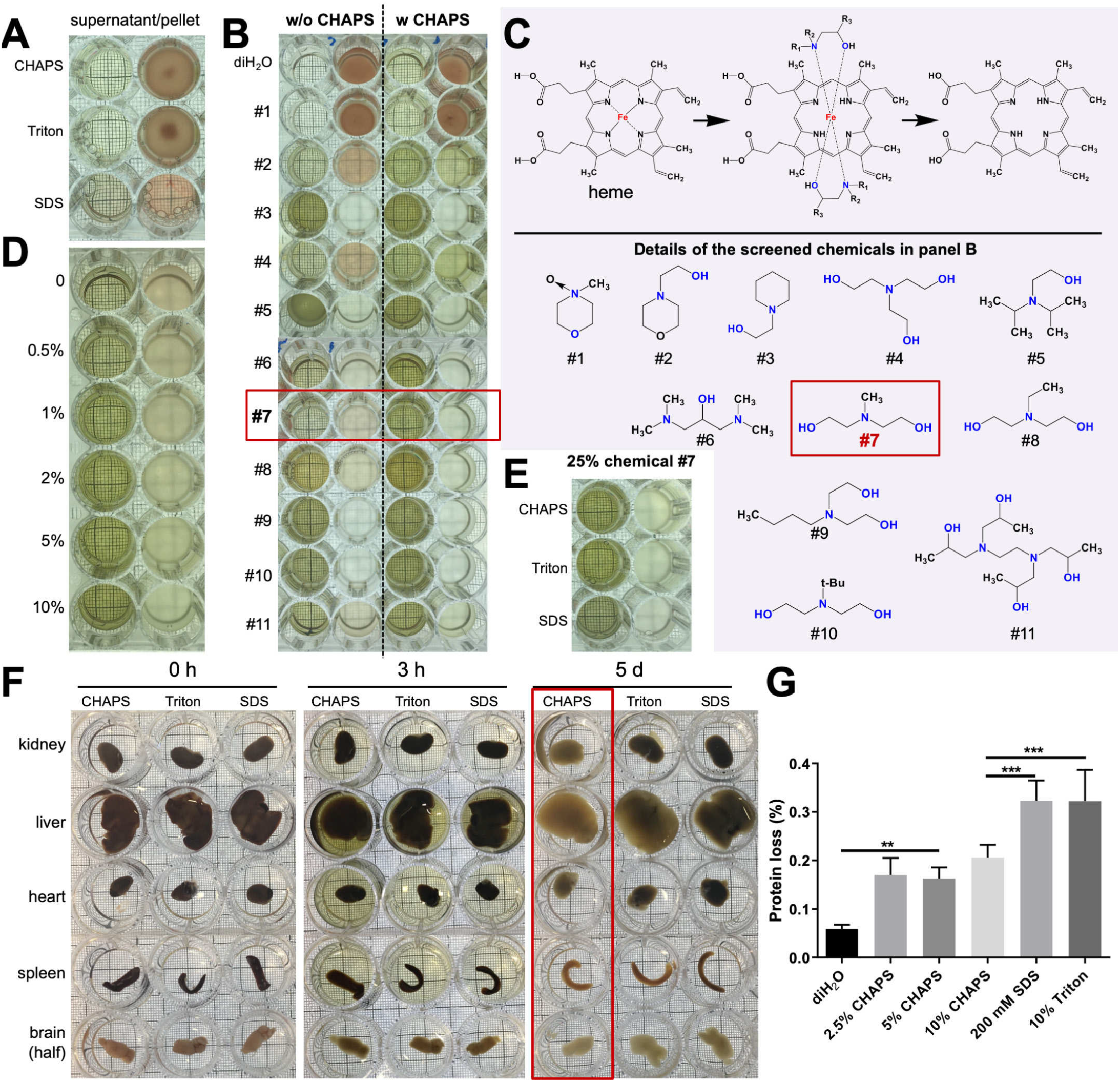
Screen for CHAPS compatible blood decolorization chemicals and protein loss assay for different detergents. **(A)** Detergents including CHAPS, Triton X-100 and SDS cannot decolorize PFA-fixed blood, showing colorless supernatant and red pellet solution. **(B-C)** Screening of 11 chemicals without or with CHAPS admixtures for blood decolorization (see **Table S2**). Good candidates show green supernatant and colorless pellet. CHAPS is compatible with most tested chemicals to improve the decolorization efficiency. **(C)** Schematic of hypothesized mechanism how chemicals interact with iron of heme for an efficient decolorization. **(D)** Optimization of CHAPS concentration combined with 25% w/v N-Methyldiethanolamine (chemical 7) for blood decolorization. **(E)** All detergents could decolorize with N-Methyldieth-anolamine. **(F)** Mouse organs were incubated with mixtures of 25% w/v N-Methyldiethanolamine and following detergents: CHAPS (10% w/v), Triton X-100 (10% w/v) or SDS (200 mM). CHAPS mixture shows superior permeabilization to remove the heme from blood-retaining mouse organs (red rectangle). **(G)** Protein loss assay indicating the superior retention of endogenous proteins after CHAPS treatment compared to other detergents (P values were calculated using one-way ANOVA test).

### Development of SHANEL Tissue Clearing

Next, we tested the accessibility of CHAPS permeabilized large mammalian organs using tissue clearing reagents. We chose to work with organic solvent-based clearing methods because they are fast and robust in addition to inducing tissue shrinkage [18], which helps to image larger pieces of intact cleared mammalian organs with the limited sample holding capacity of standard light-sheet microscopy. First, we cleared the intact brain of a 2-year-old adult pig, which was fixed with PFA by passive immersion after dissection, using standard organic solvent clearing reagents [33, 34]. We found that a combination of ethanol for dehydration, dichloromethane (DCM) for delipidation, and benzyl alcohol + benzyl benzoate (BABB) for refractive index (RI) matching was highly effective in rendering the centimeters-thick pig brain transparent after CHAPS/NMDEA permeabilization and decolorization (all together represents the SHANEL clearing) (**Fig. 3A-C**). SHANEL clearing provided in rapid transparency of ~12.0 × 7.3 × 5.0 cm size pig brain including heavily myelinated white matter, thalamus and brainstem within 1.5 months (**Fig. 3C**). The dimensions of the pig brain after clearing became 7.5 × 5.0 × 3.3 cm, with a shrinkage ratio of 67% in volume.

**Fig. 3.**
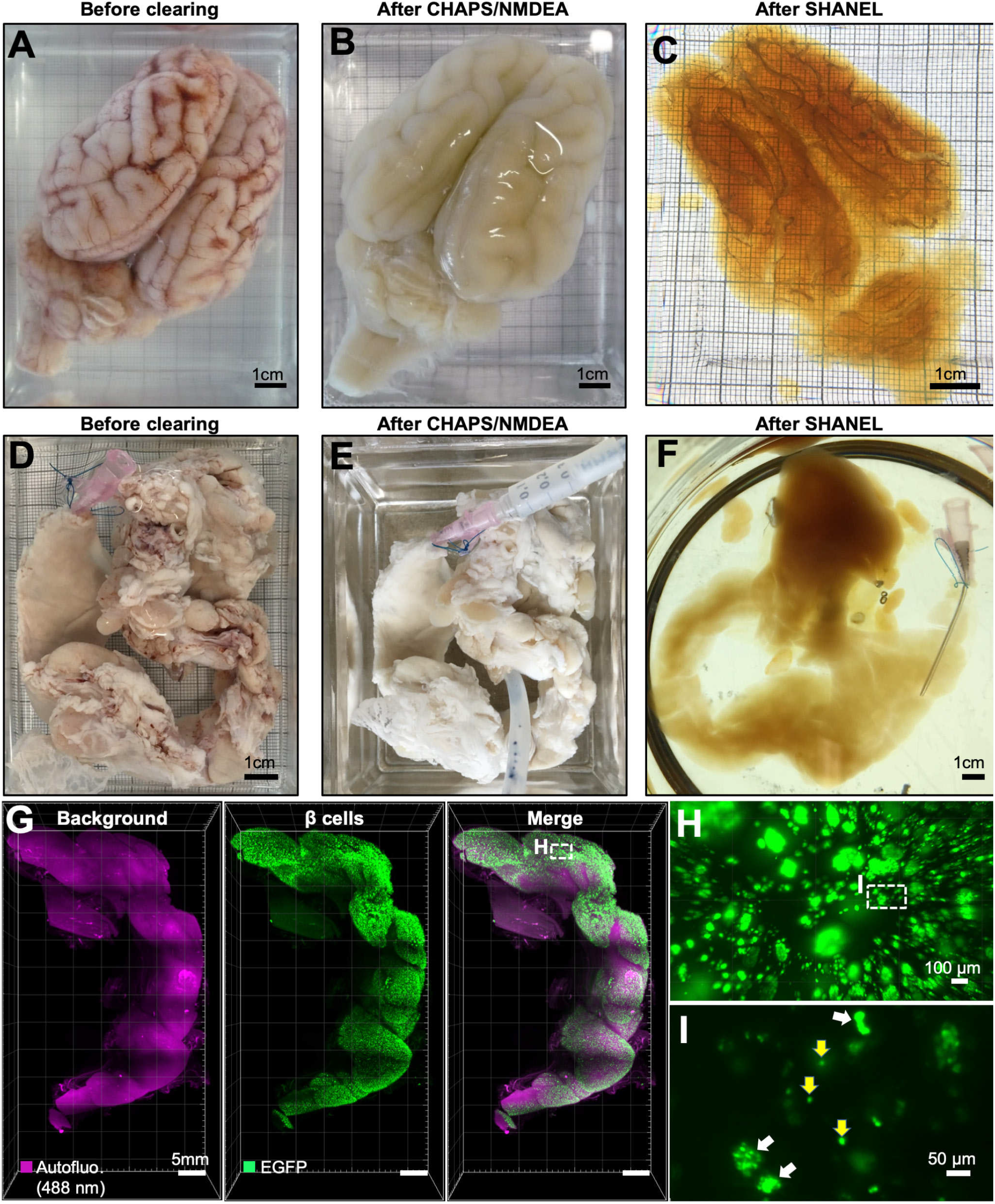
SHANEL clearing of brain and pig pancreas from adult pig. **(A)** PFA-fixed adult pig brain with retained blood. **(B)** Permeabilized and decolorized pig brain by CHAPS/NMDEA. **(C)** Fully transparent adult pig brain after SHANEL clearing. **(D)** PFA-fixed, dissected (INS-EGFP transgenic pig) pancreas with retained blood. **(E)** After CHAPS/NMDEA treatment, the pancreas was completely decolorized. **(F)** Transparent pig pancreas after SHANEL clearing. **(G)** 3D distribution of β-cell islets imaged by light-sheet microscopy after nanobody boosting of EGFP. **(H)** High magnification view of the region marked in G. **(I)** High magnification view of the region marked in H showing β-cell islets of single cells (yellow arrows) or multiple of cells (white arrows). We observed that the majority of larger islet shapes were circular or oval. See also **Supporting Fig. 2** and Supporting Movie 1.

Recent developments in gene editing with CRISPR/ Cas9 technology have enabled easy generation of large transgenic reporter mammals, expressing fluorescent proteins in the tissues of interest [35, 36]. Therefore, we applied SHANEL clearing to *INS-EGFP* transgenic pig pancreas exhibiting porcine insulin gene *(INS)* promotor driven beta cell specific enhanced green fluorescent protein (EGFP) expression in the islets of Langerhans [37] (**Fig. 3D-F**). To enhance the signal of EGFP in centimeters sized tissue, we used anti-GFP nanobodies conjugated with bright Atto dyes [14]. We demonstrated that the 3D distribution of pancreatic beta cells as single cells or groups of cells within the islets of Langerhans could be readily assessed by our new approach enabling quantification of islet volume and demonstration of islet size heterogeneity (**Fig. 3G-I; Supporting Fig. 2; Supporting Movie 1**).

### Generation of intact transparent human brain by SHANEL clearing

Labeling and clearing of the intact human brain would be a tremendous step forward to map it in the near future. As the human brain vascular system is an established network reaching all parts of the brain, we used it to deliver the chemical cocktails deep into the brain tissue (**Fig. 4A**). We used the two main pairs of large arteries, the right and left internal carotids (CR, CL) and the right and left vertebral arteries (VR, VL) to circulate solutions. First, we used PBS/heparin solution to wash out the liquid blood, followed by 4% PFA/PBS solution to fix the brain. Subsequently we isolated the whole human brain with these major vessels and connected eyes (with a volume of ~1344 cm^3^ and dimensions of ~15.0 × 10.4 × 14.4 cm) from the skull. Then we set up a pressure-driven pumping system to circulate cell nucleus labeling dye (TO-PRO-3) and all clearing reagents through the four arteries to accelerate the process (**Fig. 4B**). By doing so, we rendered the whole adult human brain transparent for the first time. To demonstrate the full transparency, we illuminated the whole brain with a condensed fluorescent light, which travelled end-to-end in the intact see-through human brain (**Fig. 4C**). This is a 2-3 orders of magnitude increase in the volume of human brain tissue that could be rendered transparent compared to prior methods [38]. The whole process takes ~3 months and costs approximately 3000 € for one adult human brain (**Supporting Table 4**). The final volume of shrunken brain was 56% of the initial volume. As there are no light-sheet microscopy systems yet available to image such large organs, we used a modified commercial light-sheet microscopy (Lavision Ultramicroscopy) to acquire mosaic images of the intact cleared human eye with a diameter of ~3 cm. We imaged TO-PRO-3 and autofluorescence signals of the intact cleared eye and reconstructed the details of its anatomical structures including sclera, iris and suspensory ligament in 3D (**Fig. 4D-E; Supporting Movie 2**). Thus, our approach provides the basis for 3D histological assessment of the whole human brain in the near future.

**Fig. 4.**
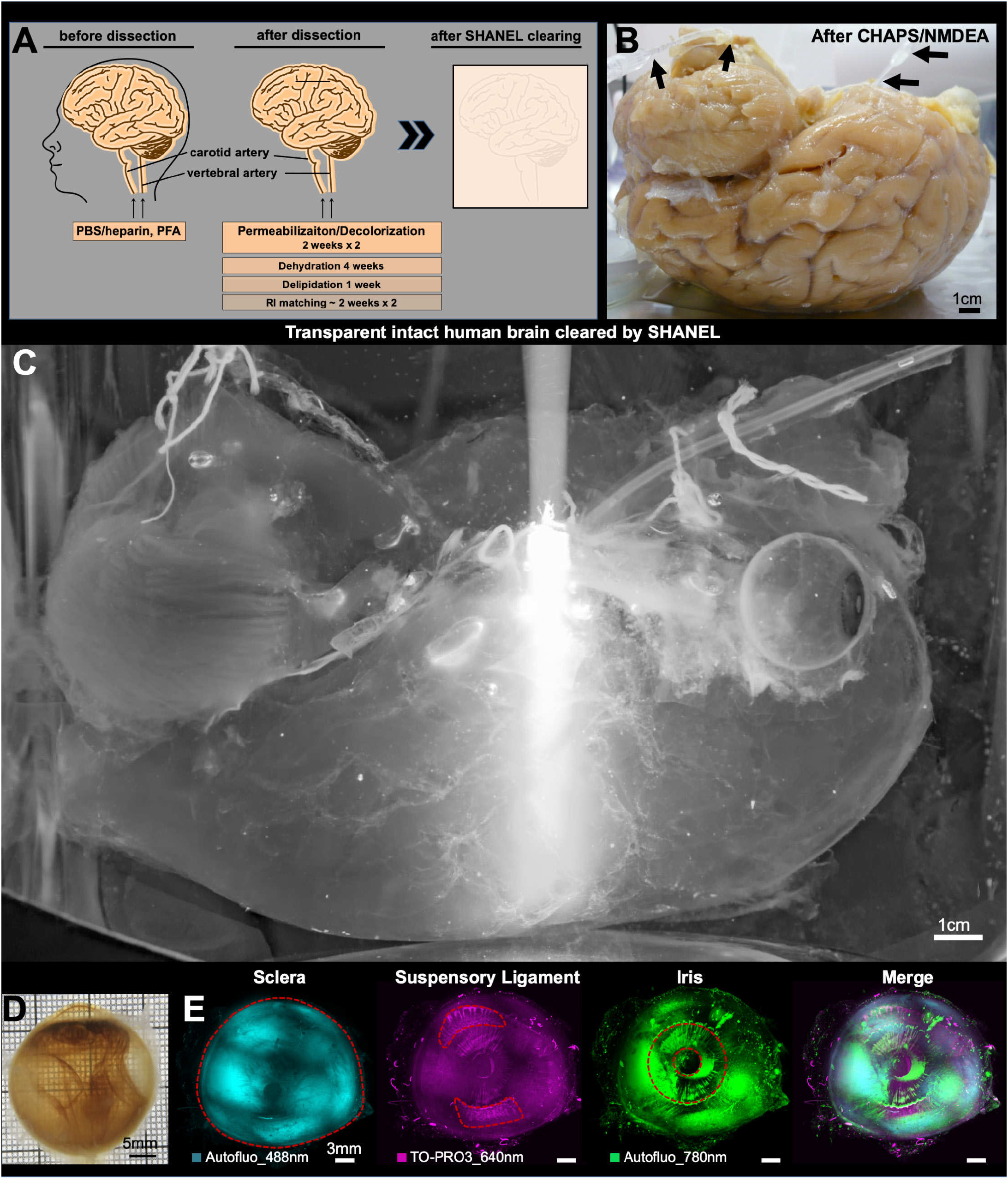
SHANEL clearing of adult intact human brain. **(A)** Active perfusion pipeline is used to accelerate whole human brain SHANEL clearing. **(B)** A sample of permeabilized and decolorized whole human brain by CHAPS/NMDEA via active pumping setup (black arrows). **(C)** Transparent whole human brain, a volume of ~1344 cm^3^ and the dimensions of ~15.0 × 10.4 × 14.4 cm. Light of epifluorescent microscope travels end-to-end demonstrating the full transparency of intact human brain. **(D)** Brightfield view of an eye, dissected from the intact transparent human brain. **(E)** 3D reconstruction of the eye from light-sheet scans showing the sclera, iris and suspensory ligament structures. See also Supporting Movie 2.

### SHANEL histology of centimeters sized human organs

Because *in vivo* genetic labeling and fluorescent dye tracing are not applicable to study post-mortem human tissue, cellular and molecular interrogation of human organs requires post-mortem staining using dyes and antibodies. However, state-of-the-art antibody labeling techniques of decades-aged human samples are limited to ≤50 μm in standard histology, and to a maximum of ~1 mm thickness in prior clearing methods, mainly due to the weak permeabilization of the tissue and slow diffusion of the antibodies, which are large molecules (~150 kDa) [11, 24, 25]. In addition, age-related accumulation of highly autofluorescent molecules increases the background tremendously in thicker tissues [39]. To overcome these limitations and achieve staining of human brain tissues as large as the current imaging systems can accommodate (1.0-1.5 cm of thick tissue) [18], we set out to develop a deep-tissue antibody labeling method for thick human tissues. Towards this goal, we perfused an intact human brain from a 92-years old female using CHAPS/NMDEA to permeabilize and decolorize. This treatment softened the sturdy human brain and allowed its easy sectioning into 12 coronal slices (each 1.5 cm thick) using a brain slicer (**Supporting Fig. 3**). The permeabilized human brain slice was further slackened by delipidation using DCM/MeOH (methanol). Next, we identified acetic acid and guanidine HCl as powerful reagents to loosen the extracellular matrix (ECM) for the diffusion of large molecules such as antibodies (**Supporting Fig. 4**).

We firstly labeled and cleared an intact 16.5 × 16.5 × 1.5 cm human brain slice using Methoxy-X04 and TO-PRO-3, which were affordable dyes in large quantities, to label Abeta plaques and cell nuclei in such large tissue, respectively. After clearing, we scanned a 7.5 × 5 × 0.4 cm region of brain slice in ~2 days using an upright confocal microscopy that is design to scan large slices of tissue. We detected Abeta plaque accumulation in several brain regions including cingulate gyrus (CG), precuneus (PCun), superior temporal gyrus (STG) and middle temporal gyrus (MTG) (**Supporting Fig. 5A-D; Supporting Movie 3**). Next, we used epifluorescence microscopy to quickly screen the 1.5 cm thick half human brain slice and again readily located the regions with Methoxy-X04-labeled Abeta accumulation for subsequent confocal microscopy imaging (**Supporting Fig. 5E-H**).

Antibody labeling of tissues has been widely used to interrogate the specific cellular architecture and underlying molecular mechanism of biological processes. Therefore, we next applied SHANEL to assess the possibility of antibody-based histology of centimeters sized human tissues. Towards this goal, we first used ionized calcium binding adaptor molecule 1 (Iba1) antibody to immunolabel microglia (**Fig. 5A-H**). Iba1 + microglia were identifiable throughout the 2.0 × 1.8 × 1.5 cm size human brain slice (**Supporting Movie 4**). We also observed morphological differences: microglia cells in gray matter were mostly larger and more ramified compared to white matter (**Fig. 5E-H**). Next, we used tyrosine hydroxylase (TH) antibody to immunolabel neuronal structures along with PI labeling of the cell nuclei (**Fig. 5I-L**). We could clearly see specifically labeled axonal extensions throughout the 1.8 × 1.8 × 1.5 cm human brain slice (**Fig. 5J-L; Supporting Movie 5**). These results demonstrate that SHANEL histology can successfully permeabilize entire 1.5 cm thick sturdy human brain slices for deep tissue antibody labeling. This represents a 1-2 orders of magnitude enhancement in the thickness of adult human tissue that could be processed for histology with prior methods [38]. Studying vasculature has been a valuable method to explore diverse developmental and pathological phenomena in biological tissues. Again, histological assessment of vasculature in human tissue has been limited by the penetration of specific dyes and antibodies. Here, we used Lectin dye to effectively label a human brain sample with a size of 3.0 × 1.9 × 1.5 cm (**Supporting Fig. 6**). In addition to specific vascular structures (e.g. **Supporting Fig. 6C**), we could identify vasculature related tissue abnormalities, seen as swollen structures in our deep tissue Lectin labeling (e.g. **Supporting Fig. 6D**). Thus, SHANEL histology allows the labeling of specific molecules, cells and vasculature in centimeters sized human brain samples presenting a viable tool to scale up the investigation of brain pathologies.

**Fig. 5.**
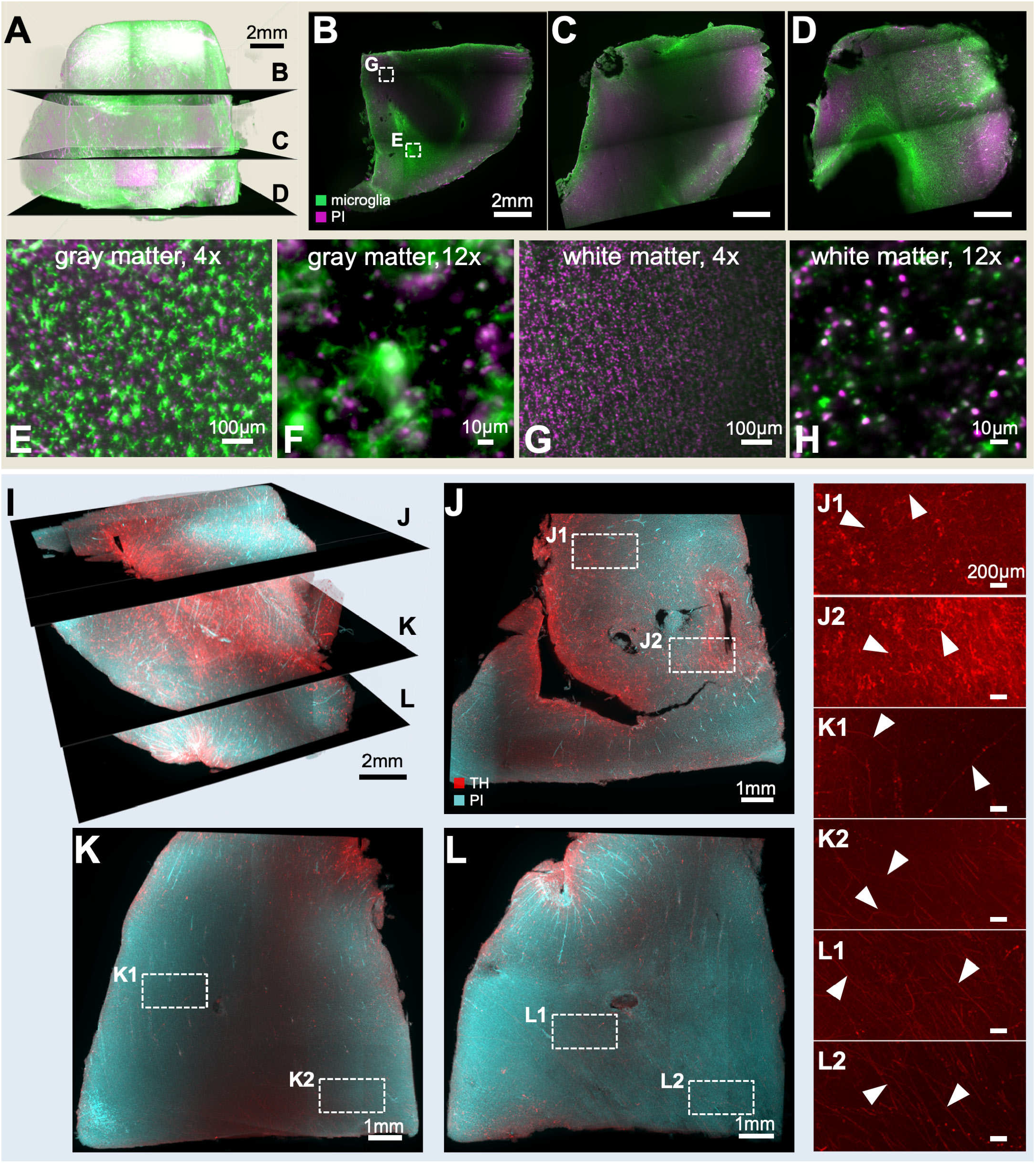
SHANEL histology on centimeters thick human tissues imaged by light-sheet microscopy. **(A-D)** Iba1 microglia (green) and propidium iodide (PI) (magenta) labeling of post-mortem human brain tissue with an original size of 2.0 × 1.8 × 1.5 cm (1.3 × 1.2 × 1.0 cm after shinkage). Individual microglia throughout the gray matter **(E, F)** and white matter **(G, H)** are evident. **(I)** Tyrosine hydroxylase (TH) (red) and propidium iodide (PI) (cyan) labeling of post-mortem human brain tissue with an original size of 1.8 × 1.8 × 1.5 cm (1.2 × 1.2 × 0.91 cm after shinkage). **(J-L)** TH+ axonal extensions in gray (white arrowheads in J1-J2) and white matter (white arrowheads in K1-L2) throughout the entire depth of tissue are evident. Note that cyan channel is not shown in J1-L2 to emphasize the TH labeling. See also Supporting Movies 4 and 5.

Next, we tested our SHANEL technology on entire human kidneys using affordable small dyes. There is a huge shortage of donor organs for hundreds of thousands of people in need of kidney transplantation [40]. The waiting time for donation may be several years, and cost of transplantation might reach about half a million dollars [40]. Understanding the 3D structure of the human kidney would be of value for tissue engineering efforts aiming to generate artificial kidneys [41], which require detailed cellular and molecular knowledge on intact human kidney tissues. Kidneys are the major organ for blood filtration through glomeruli, whose density and size are critical for healthy organ function. Towards understanding the 3D cellular structure of the human kidney we used SHANEL histology with active perfusion of TRITC-dextran and TO-PRO-3 dyes through the renal artery to label the vessels and dense cellular structure of the glomeruli in the entire kidney with a size of 11.5 × 8.2 × 3.0 cm. After labeling, we also actively pumped the clearing reagents through the kidney to overcome the age and size related challenges. We achieved full transparency, revealing the primary renal artery, secondary branches of segmental arteries and interlobar arteries (**Fig. 6A-C**). Using light sheet microscopy, we could visualize the 3D distribution of vessels and glomeruli in the kidney cortex over large volumes (1.2 × 1.2 × 0.45 cm), and trace individual afferent arteriole and its glomeruli (**Fig. 6D-F; Supporting Movie 6**). Through cortex profile counting, we found that the width of the cortex zone was around 2742 ± 665 μm (mean ± s.d.), the diameter of glomerular caliper was 221± 37 μm, and afferent arteriole diameter was 71 ± 28 μm (**Fig. 6G**). Thus, our technology provides accurate and precise mapping of glomeruli and arteries of kidney in 3D at a reasonable time and cost.

**Fig. 6.**
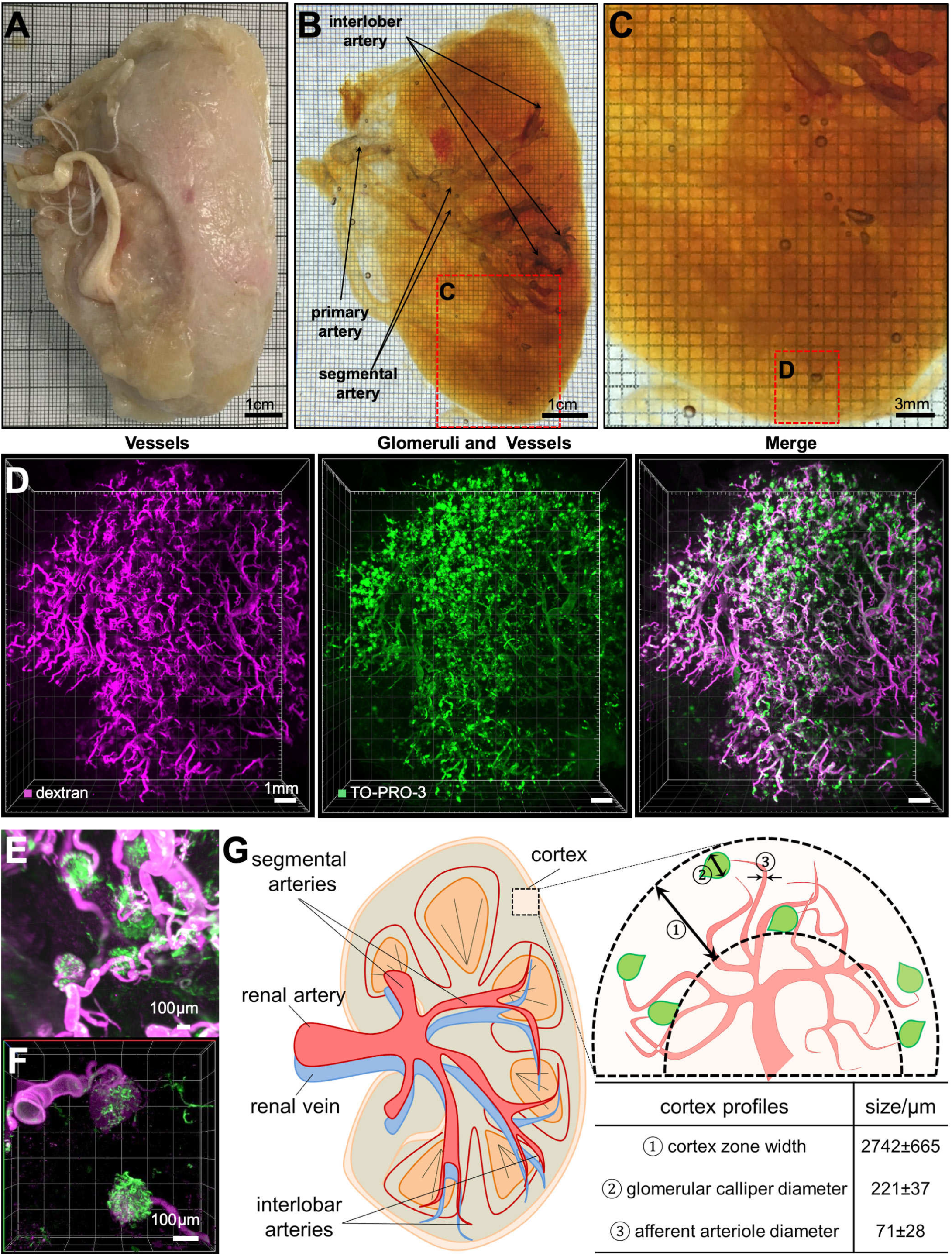
Cellular investigation of human kidney. **(A-C)** A PFA-fixed adult human kidney **(A)** was rendered totally transparent after SHANEL clearing, revealing visible arteries **(B,C). (D)** 3D reconstruction of vessels and glomeruli of the kidney cortex region marked in (C) imaged by light-sheet microscopy. TRITC-dextran labels the vessels (magenta), while TO-PRO-3 labels glomeruli more prominently (green). High magnification light-sheet microscopy **(E)** and confocal microscopy **(F)** images show the structural details of afferent arteriole (magenta) and glomeruli (green). **(G)** Human kidney anatomy and cortex profiles from the 3D reconstruction. See also Supporting Movie 6.

## DISCUSSION

Histological studies of human tissues suffer from a lack of scalable methods to label and image large human specimens. Here we present the SHANEL method, which is derived from a new tissue chemistry achieving thorough permeabilization of fixed post-mortem human tissues. It allows full penetration of labeling molecules and clearing chemicals into centimeters sized decades-aged human tissues such as post-mortem human brain and kidney specimens. Our method can also readily be applied to many organs in parallel as it does not require lengthy handwork other than setting up the perfusion system and exchange of the solutions. Thus, this scalable method could vastly accelerate the 3D structural and molecular mapping of cells in intact human organs including the human brain.

Early efforts of adult human organ clearing started a century ago with slow progress in transparency and labeling options [42]. A particular difficulty has been the age dependent accumulation of intracellular and extracellular molecules such as lipofuscin and neuromelanin pigments. Lipofuscin is a mixture of highly oxidized cross-linked macromolecules including proteins, lipids, and sugars from different cellular metabolic processes [27]. Age-related accumulation of lipofuscin in the human body, in particular in the brain, correlates with senescence and sturdiness of human tissues [27]. Similarly, insolubility of the collagen also increases in the human body with the ageing, leading to hardening, browning and autofluorescence of the tissues [28]. Due to age-related accumulation of such insoluble macromolecules in human tissues over several decades, histological examination relying on the penetration of large molecules such as antibodies deep into tissues has been very challenging. Recently developed tissue clearing methods have proven to be a promising way to achieve histological assessment of intact specimen. While diverse clearing methods have been quite successful in rodents (with the age of a few months), they have not been as effective on decades-aged human tissues. We solved this problem by developing a new method to permeabilize the sturdy human tissues, which is the prerequisite step for any tissue labeling and clearing method.

Here, we hypothesize that the micellar structures of detergents would be critical to penetrate the dense meshes of sturdy human tissues. Towards this goal, we identified CHAPS, a zwitter-ionic detergent having rigid steroidal structure with hydrophobic and hydrophilic faces. The ‘facial’ amphiphilicity of CHAPS is responsible for the aggregation behavior and surface configuration of molecules at interface. Firstly, CHAPS aggregates into much smaller micelles compared to standard ‘head-to-tail’ detergents such as SDS and Triton X-100, facilitating its rapid penetration deep into dense human tissues. In aggregation formation, the hydrophobic faces (β-plane of steroid) are considered to contact with each other, while the hydrophilic ones (three hydroxyl groups) remain exposing to the aqueous environment. Secondly, in terms of interfacial interaction, the facial amphiphiles bind to the bilayer in a unique mode that allows coverage of a larger hydrophobic area instead of being embedded into the bilayer. By this way, less facial amphiphile CHAPS is needed and no residual CHAPS left behind in the tissue after the treatment. Moreover, as a mild zwitterionic detergent, CHAPS exhibits better preservation of tissue endogenous biomolecules compared to SDS and Triton X-100 for cellular and molecular interrogation. Starting from this new permeabilization chemistry, we further developed our technology by loosening ECM using acetic acid and guanidine HCl [43, 44].

The resulting SHANEL histology method enabled diffusion of molecules as large as conventional IgG antibodies for straightforward 3D histology of centimeters sized human organs. Thus, our approach can also tremendously help scaling up the efforts on Human Protein Atlas (HPA) by reducing the time to label and annotate across large human tissues [45, 46]. Yet, our method does not fully eliminate the tissue autofluorescence, in particular those coming from the lipofuscin in the brain tissue. However, this autofluorescence can be used to extract more information on senescence of cells throughout different tissue layers in addition to imaging of specific dye/ antibody signals in other channels.

The SHANEL technology is also applicable to other large mammal organs. As pig is a much better model system for the human islets research compared to rodents, study of transgenic *INS-EGFP* pig pancreas in combination with SHANEL clearing can also accelerate research in metabolic disorders. We demonstrate that the 3D distribution of pancreatic beta cells as single cell or groups of cells within the islet of Langerhans can be readily imaged and quantified by our new approach demonstrating the islet size heterogeneity. Regional differences of islet size and distribution (head vs. tail) in human pancreas are well known, and alterations in beta cell mass occur in diverse metabolic disorders [47]. For instance in disease conditions like type 2 diabetes, regional changes of islet distribution leading to preferentially large islets in the head region occur [48]. Another interesting application would be to assess the quantity and distribution of porcine islets after intraportal xenotransplantation into the liver of non-human primate models [48, 49].

Currently, we lack fluorescent microscopy systems that can image biological specimen as large as the intact human brain. While we rendered the entire human brain transparent, we could not use current laser-scanning light-sheet microscopes to construct a complete 3D image of the whole brain. The development of such imaging systems with extended stages and imaging capacities would tremendously accelerate studies on phenotyping of the cellular and molecular architecture of the whole human brain. Leading towards this possiblity, Voigt et al., recently proposed the mesoSPIM (mesoscale selective plane illumination microscopy), an open-hardware microscopy platform for imaging cleared tissues several centimeters in size [50]. Here, we sliced the intact adult human brain into twelve 1.5 cm-thick sections using standard slicer equipment. Still, such thick slices are much easier to handle compared to micrometer or even millimeter thick sections, because there is no need to embed or collect them on glass slides. Moreover, SHANEL histology allowed a thorough labeling of these 1.5 cm-thick brain slices. Because we use organic solvents for tissue clearing, we also reduced the thickness of the cleared samples by about one third. This allows the usage of high numerical aperture objectives such as 4x Olympus objective (NA: 0.28 and working distance (WD): 10 mm) and 20x Zeiss Clarity objective (NA: 1, WD: 5.6 mm) on defined regions of interest to reconstruct the entire 1.2 – 2 cm thick human tissue by scanning from both sides. Another advantage of our approach is that, after SHANEL histology, the specimens become hard, enabling easy handling of complete organs and slices. Finally, owing to the complete dehydration and incubation in organic solvents, the tissues are preserved by SHANEL histology, allowing long-term storage for the future studies by the same investigators or in other labs. This approach will reduce the need of repetitive sample collection to perform similar experiments with the same type of post-mortem human tissues.

In conclusion, this work presents a new technology to permeabilize centimeters sized aged-human tissues for molecular and cellular phenotyping. This method allows deep tissue labeling and clearing of human specimens as large as the intact human kidney and brain. Thus, SHANEL histology can be a key technology to map intact human organs in the near future, which would accelerate our understanding of the physiological and pathological conditions governing human life.

## EXPERIMENTAL PROCEDURES

### Small-angle X-ray scattering measurements

We used small-angle X-ray scattering to determine the size, shape, and aggregation number of CHAPS and SDS micelles. Experimental data were collected at beam line 12ID at the Advanced Photon Source (APS) using procedures as previously described [51–53]. In brief, measurements were carried out with a custom-made sample cell and holder [54], at a temperature of 25 °C and an X-ray energy of 12 keV, with a sample-to-detector distance of 1.8 m. We defined the magnitude of momentum transfer as *q = 4π/λ · sin (θ*), where *2θ* is the total scattering angle and *λ* = 1 Å the X-ray wavelength. The useable q-range in our measurements was 0.02 Å^−1^ < *q* < 0.275 Å^−1^. Scattering angles were calibrated using a silver behenate standard sample and data read out, normalization, and circular averaging were performed using custom routines at beam line 12ID, APS. SDS and CHAPS were measured in PBS buffer for 10 exposures of 0.1 s. Matching buffer profiles were subtracted for background correction. Subsequent exposures were compared to verify the absence of radiation damage. Horse heart cytochrome *c* at 8 mg/ml was used as a molecular mass standard.

### Small-angle X-ray scattering data analysis

Radii of gyration *R_g_* and forward scattering intensities *I(0)* were determined from Guinier analysis [55–57] of the low *q* region of scattering profiles, i.e. from a fit of the logarithm of the scattering intensity vs. *q*^2^ for small *q* (**Supporting Fig. 1A,B**). The fitted radii of gyration are in excellent agreement with previously reported values, as far as available (**Fig. 1C**). In addition, the full scattering profiles were fitted with one- and two-component ellipsoid models [51] (**Supporting Fig. 1C**). We found that the CHAPS data were well described by a prolate one-component ellipsoid model with long axis of ≈30 Å and short axis ≈12 Å. In contrast, the SDS data require a two-component ellipsoid model for a convincing fit, similar to other ‘head-to-tail’ detergents [51], and were well described by a oblate two-component ellipsoid model with an inner core (representing the region occupied by the hydrophobic tail groups) with small axis ≈15 Å and long axis ≈25 Å (in good agreement with results from neutron scattering [58]), surrounded by a shell (representing the head groups) of thickness ≈3.5 Å.

Aggregation numbers for CHAPS and SDS micelles were determined using two independent approaches [51]. One approach was to use the fitted geometric models to compute the volumes of the micelle (in the case of CHAPS) or of the hydrophobic core (in the case of SDS) and to compute the aggregation number by dividing the total volume by the volume of the monomer (for CHAPS, see **Table S1**) or the hydrophobic core volume by the volume of the alkyl tail (determined from the Tanford formula [59], 350.2 Å^3^ for SDS). We note that the approach of using the micelle volume to determine the aggregation number uses the entire scattering profile in the model fit, but is independent of the scale of the scattering intensity, since only the shape of the scattering pattern is fit. An alternative and independent approach to determining the aggregation number is to determine the forward scattering intensity *I(0)* from Guinier analysis and to compute the aggregation number from the equation [51]:

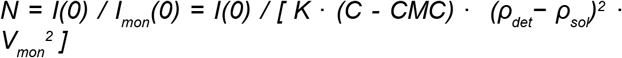

Here *K* is a proportionality constant that was determined from measurements of horse heart cytochrome *c* as a scattering standard, *C* is the detergent concentration, *CMC* the critical micelle concentration, *ρ_det_* the electron density of the detergent, *ρ_sol_* = 0.34 e/Å^3^ the electron density of the solvent, and *V_mon_* the detergent monomer volume. Values for *CMC*, *ρ_det_* and *V_mon_* are given in **Table S1**. The approach of using *I(0)* to determine the aggregation number only uses the very low q information in the scattering pattern and is independent of any model assumptions about the size or shape of the micelles. Aggregation numbers determined by the independent approaches are in good agreement.

### Mouse samples

CD-1 IGS (Charles River, stain code: 022) mice were used for blood collection, organs screening and protein loss assay. The animals were housed under a 12/12 hours light/dark cycle. The animal experiments were conducted according to institutional guidelines: Klinikum der Universität München / Ludwig Maximilian University of Munich and after approval of the Ethical Review Board of the Government of Upper Bavaria (Regierung von Oberbayern, Munich, Germany) and in accordance with the European directive 2010/63/EU for animal research. All data are reported according to the ARRIVE criteria. Animals were randomly selected. Mice were deeply anesthetized using a combination of medetomidine, midazolam and fentanyl (MMF; 5 mg, 0.5 mg and 0.05 mg per kg body mass for mice; intraperitoneal). As soon as the animals did not show any pedal reflex, blood was extracted intracardially from the left ventricle. Then the blood was mixed with 2 times volume of 4% paraformaldehyde (PFA) in 0.01 M PBS (pH 7.4; Morphisto, 11762.01000) and incubated for 24 h at 4 °C. Animals for protein loss assay were intracardially perfused with heparinized 0.01 M PBS (10 U ml-1 of heparin, Ratiopharm; ~110 mmHg pressure using a Leica Perfusion One system) for 5–10 min at room temperature until the blood was washed out, followed by 4% PFA for 10–20 min. The brains were dissected and post-fixed in 4% PFA for 1 d at 4 °C and later washed with 0.01 M PBS for 10 min 3 times at room temperature.

### Pig samples

Pig intact brain was brought from local slaughterhouse and immersion fixed for 10 days with 4% PFA at 4°C. Housing, breeding and animal experiments of INS-EGFP transgenic pigs were done at the Institute of Molecular Animal Breeding and Biotechnology at LMU Munich according the approval of the Ethical Review Board of the Government of Upper Bavaria (Regierung von Oberbayern, Munich, Germany) and in accordance with the European directive 2010/63/EU for animal research. All data are reported according to the ARRIVE criteria. INS-EGFP transgenic pigs have a beta-cell specific EGFP reporter gene expression driven by the porcine insulin *(INS)* promoter [37]. Pancreas of an exsanguinated 5.5 month old INS-EGFP transgenic male pig of a German landrace background was dissected, its pancreatic ducts was cannulated and 20 ml of ice-cold 4% PFA was slowly injected.

### Human samples

Intact human brains were taken from different human body donors with no known neuropathological diseases. All donors gave their informed and written consent to explore their cadavers for research and educational purposes, when still alive and well. The signed consents are kept at the Anatomy Institute, University of Leipzig, Germany. Institutional approval was obtained in accordance to the Saxonian Death and Funeral Act of 1994. The signed body donor consents are available on request.

The brains of a 92 years old female and a 48 years old male of the body donation program of the Institute of Anatomy, University of Leipzig were fixed in situ by whole head perfusion via carotid and vertebral arteries under a pressure of below 1 bar. The head was firstly perfused with 5 L heparinized 0.01 M PBS (10 U ml-1 of heparin, Ratiopharm), followed by 3 L 4% PFA in 0.01 M PBS for 2-3 h. The veins were finally closed to maintain the PFA to the brain. Then the brains were recovered by calvarian dissection and preserved at least 1-2 weeks for postfixation submersed in the 4% PFA solution. The kidneys of a 93 years old female were dissected from the body. The blood was flushed by 200 ml of heparinized PBS in a PBS bath for 1 h and perfused with 400 ml of PFA immersed in PFA solution. The kidneys were preserved at least 1-2 weeks for post-fixation submersed in the 4 % PFA solution at 4 °C.

### Screening of affordable and scalable chemicals for blood decolorization

PFA-fixed mouse blood was thoroughly vortexed with only 25% w/v screened chemicals (Sigma, **Table S2**) or only detergents (10% w/v CHAPS, 10% w/v Triton X-100 or 200mM SDS, Sigma), or the mixture of detergents and 25% w/v chemicals, and then immediately centrifuged at 15000 rpm for 5 min at room temperature. Supernatant was transferred into the multi-well plate and pellet was dissolved in diH_2_O and also transferred into paired well. Bright-field images were immediately captured. In accordance with our hypothesis that electron-rich nitrogen donor and polarizable hydrogen of chemicals tend to bind with iron of heme as double-action multidentate ligands, then eluting the heme from the blood, all of the tested chemicals could partly decolorize the red heme. These effects were improved with the addition of detergents. Considering the practical prices and availability of reagents to process large samples such as the human brain, we optimized the concentration of CHAPS with N-Methyldiethanolamine (chemical 7) for later experiments [17].

### Permeabilization and decolorization of mouse organs with remaining blood

Mouse organs were dissected and post-fixated with 4% PFA/PBS as soon as the animals did not show any pedal reflex. The blood-retaining organs were washed with PBS for 3h × 3 times and put into a multi-well plate and bright-field images were captured. Then the organs were incubated with mixtures (10% w/v CHAPS and 25% w/v N-Methyldiethanolamine, 10% w/v Triton X-100 and 25% w/v N-Methyldiethanolamine, 200 mM SDS and 25% w/v N-Methyldiethanolamine) at 37 °C on a shaking rocker (IKA, 2D digital). The solutions were refreshed when the color changed to green until colorless. Then the grouped samples were washed with PBS 3 times for 3h at room temperature and imaged again under bright-field.

### Protein loss assay

PFA-fixed adult mouse (3-4 month) brains were cut into 1-mm-thick sections using a vibratome (Leica, VT1200S) [11, 22]. All sections were weighted and randomly grouped then placed in 5 ml solutions as follow: distilled water, 2.5% w/v CHAPS, 5% w/v CHAPS, 10% w/v CHAPS, 200nM SDS or 10% w/v Triton X-100. The samples were incubated for 2 weeks at 37 °C on a shaking rocker. The respective solutions and quantity of protein lost from tissue by diffusing into solutions was measured using Bio-Rad DC protein assay kit (bio-rad, 5000116). Total protein in mouse brain was estimated at 10% (wt). For each group, the standard solution was prepared in the same buffer as the sample.

### Passive SHANEL clearing of pig brain

PFA-fixed pig brain samples were washed with PBS at room temperature, incubated with 10% w/v CHAPS and 25% w/v N-Methyldiethanolamine solution at 37 °C on a shaking rocker. The incubation time of whole brain was 10 days with solution refreshed once at day 5. After PBS washing at room temperature, the samples were shaken with a series of Ethanol/DiH_2_O solutions (50%, 70%, 100%, 100% v/v) at room temperature, following further DCM (Roth, KK47.1) incubation, immersed into BABB solution until complete transparency. The incubation time depended on the sample size, for whole brain, 4 days for each step.

### Active SHANEL clearing of pig pancreas

Pancreas from a 5.5-month old INS-EGFP transgenic pig was dissected without the tail part, a G 20 venous catheter was inserted into the pancreatic duct and sewed on. 4% PFA was injected to fix the tissue, followed by post-fixation for 3 days at 4 °C. Then the sample was set up in the active pumping system consisting of a peristaltic pump (ISMATEC, REGLO Digital MS-4/8 ISM 834), chemical-resistant PTFE tubing (VWR, 228-0735) and a glass chamber. After PBS washing, 200 ml solution of 5% w/v CHAPS and 12.5% w/v N-Methyldiethanolamine was circulated through the pancreas for 8 days in total, refreshing the solution with fresh one every 2 days. After 2 times of PBS washing for 3 h, the sample was pretreated with 200 ml of permeabilization solution containing 1.5% goat serum, 0.5% Triton X-100, 0.5 mM of methyl-β-cyclodextrin, 0.2% trans-1-acetyl-4-hydroxy-l-proline and 0.05% sodium azide in PBS for half a day at room temperature. Subsequently, the perfusion proceeded further, through connection of a 0.20 μm syringe filter (Sartorius, 16532) to the intak-ending of the tube to prevent accumulation of dye aggregates into the sample. At the same time, an infrared lamp (Beuer, IL21) was used to heat up the solution to 26–28 °C. With this setting, the pancreas was perfused for 13 d with 250 ml of the same permeabilization solution containing 30 μl of Atto647N-conjugated anti-GFP nanobooster (Chromotek, gba647n-100). After that, the pancreas was washed out by perfusing with washing solution (1.5% goat serum, 0.5% Triton X-100, 0.05% of sodium azide in PBS) for 6 h 3 times at room temperature and PBS for 3 h at room temperature. The clearing was started with series of 250 ml of EtOH/DiH_2_O solution (50%, 70%, 100%, 100% v/v) pumping for 6 h at each step. Then the pancreas was passively incubated with DCM for 1 day, proceeded by BABB solution until complete transparency in one month.

### Active SHANEL clearing of intact human brain

The four main arteries of PFA-fixed intact human brain were connected to a peristaltic pump (ISMATEC, REGLO Digital MS-4/8 ISM 834) through a chemical-resistant PTFE tubing (VWR, 228-0735) in a glass chamber. 5 L of 10% w/v CHAPS and 25% w/v N-Methyldiethanolamine solution was pumped continuously into arteries keeping the pressure at 180-230 mmHg (50-60 rpm). One channel from the pump, made by a single reference tube, was set for circulation of the solution through the artery into the brain vasculature system: one ending of the tube was connected to the tip which inserted into the artery tubing, and the other ending was immersed in the solution chamber where the brain was placed. The perfusion tip pumped appropriate solution into the artery, and the other ending collected the solution inside of the brain to recirculate the solution, pumping it back into the brain. At the same time, the solution was also stirred using a blender (IKA, RCT B S000) and heated to 37–39 °C. With this setting, the human brain was perfused for one month with once refresh at day 15. Then the solution was changed to PBS for washing 2 days. Using the same setting without heating, the human brain was labeled with TO-PRO-3 in 2 L PBS (1:2000 dilutions) for 1 month at room temperature. After labeling the clearing was performed by perfusing with 5L of the following series of EtOH/DiH_2_O solutions: 50%, 70%, 100%, 100% v/v for one week for each step, followed by perfusion of 5 L of DCM for another week to delipidate, in the end the sample was perfused with 5 L of BABB solution until complete transparency. When the brain was getting transparent, the BABB was refreshed and the brain was stored in this solution at room temperature without further circulation or stirring.

### 1.5 cm-thick human brain slice preparation

PFA-fixed intact human brain was actively pumped with 5 L 10% w/v CHAPS and 25% w/v N-Methyldiethanolamine solution for one month with once refresh at day 15. Then the solution was changed to PBS for actively washing for 2 days. The intact human brain was cooled in PBS at 4°C overnight, then directly cut into 1.5 cm-thick slices in coronal plane using a Rotation Cutting Slicer (Rotation Schneidemaschine, Biodur, Heidelberg, Germany). The total 12 slices were serially labeled and stored in 70% EtOH at 4°C.

### Passive histology of 1.5 cm-thick intact human brain slice

1.5 cm-thick intact human brain slice (Number 7, see Fig. S3) was randomly chosen and passively incubated with 400 ml TO-PRO-3 (T3605, Thermo Fisher, 1:2000 dilution) in PBS at room temperature for 1 week. Then the solution was changed to 400 ml with 100 μM of Methoxy X-04 (4920, Tocris) in 40% EtOH (pH=10 adjusting by NaOH solution) for another week. After labeling, the slice was washed with PBS for 1 day. The clearing started with dehydration using a series of 1 L of EtOH/DiH_2_O solutions (50%, 70%, 100%, 100% v/v) and followed by delipidation using 1 L DCM, all steps lasted 1 day. 1 L BABB solution was replaced and incubated at room temperature until completely transparency.

### Passive SHANEL antibody histology study of 1.5 cm-thick human brain samples

1.5 cm-thick intact human brain slices (Number 4 and 6, see Fig. S3) were randomly chosen and dehydrated with series of 1 L EtOH/DiH_2_O (50%, 70%, 100%, 100% v/v), then delipidated with 2 L DCM/MeOH (2:1 v/v), then rehydrated with series of 1 L EtOH/DiH_2_O (100%, 70%, 50%, 0% v/v) at room temperature. After incubating with 1 L 0.5 M acetic acid in DiH_2_O, the solution was changed to mixture of 4 M guanidine HCl, 0.05 M sodium acetate and 2% Triton X-100, pH=6.0, at room temperature to loosen the extracellular matrix. The incubation time for each solution was 1 day. Next, the slice was shortly incubated with 10% w/v CHAPS and 25% w/v N-Methyldiethanolamine solution for 4 h and washed with PBS for 1 day. The intact slice was stored in blocking buffer (0.2% Triton X-100/10% DMSO/10% goat serum/ PBS/ 0.05% of sodium azide) at 4 °C. Interested pieces were cut and incubated with the same blocking buffer at 37 °C for 1 day. Then the samples were incubated with rabbit antibody anti-Iba1 (1:1000, 019-19741, Wako), rabbit antibody anti-TH (1:250, ab112, abcam) in antibody incubation buffer (3% goat serum/3% DMSO/0.2%Tween-20/10mg·L-1 Heparin/PBS) for 1 week at 37 °C. After the primary antibody incubation, samples were washed with washing buffer (0.2%Tween-20/10mg·L-1Heparin/ PBS) for 1 day with 3 times refresh and incubated with dye-conjugated secondary antibodies (1:500, A-21245, Thermo Fisher) in antibody incubation buffer for 1 week at 37 °C. Also, samples were incubated with DyLight 649-lectin (1:500, DL-1178, Vector) in antibody incubation buffer for 1 week at 37 °C. After washing with PBS, propidium iodide (1:100, P3566, Thermo Fisher) or TO-PRO-3 dye was added in PBS for cell nuclei staining for 3 days. After labeling, the samples were dehydrated by solutions of EtOH/DiH_2_O (50%, 70%, 100%, 100% v/v) and delipidated by DCM solution for 4 h each solution. BABB solution was replaced and incubated at room temperature until completely transparent.

### Active SHANEL histology study of intact human kidney

The PFA-fixed intact kidneys were pumped through primary artery with 36 ml of mixture of 25 mg/ml tetramethylrhodamine isothiocyabate-dextran, 2 mM p-maleimidophenyl isocyanate (PMPI) and 5 mM DL-dithiothreitol (DTT). Then the kidneys were sealed into plastic bag and incubated in 37 °C overnight. After the vessel labeling, we set up the perfusion system with peristaltic pump and PTFE tubing at room temperature as human brain. The first step consisted of the washing with 2 L PBS washing for one day twice. The second step was decolorization and permeabilization with 2 L 10% w/v CHAPS and 25% w/v N-Methyldiethanolamine solution for one week. The third step was dehydration with series of 2 L EtOH/DiH_2_O (50%, 70%, 100% v/v) 4 days each solution. The fourth step was delipidation with 4 L DCM/MeOH (2:1 v/v) sealed for 4 days. The fifth step was rehydration with series of 2 L EtOH/DiH_2_O (100%, 70%, 50%, 0% v/v) 4 days each solution. The sixth step was loosening ECM with 0.5 M acetic acid for 4 days and mixture of 4 M guanidine HCl, 0.05 M sodium acetate and 2% Triton X-100, pH=6.0, for 4 days. The seventh step was again PBS washing one day twice. Next, 1 L of TO-PRO-3 dye (1:2000 diluted) was continually pumped for 2 weeks, then washed with PBS one day twice. The clearing was started with series of 2 L of EtOH/DiH_2_O solutions (50%, 70%, 100%, 100% v/v) pumping for 2 days of each step, followed by 2 L of DCM delipidation for 3 days, then proceeded to 2 L BABB solution until complete transparent.

### Light-sheet microscopy imaging

Single plane illuminated (light-sheet) image stacks were acquired using Ultramicroscopy (LaVision BioTec), featuring an axial resolution of 4 μm with following filter sets: ex 470/40 nm, em 535/50 nm; ex 545/25 nm, em 605/70 nm; ex 580/25 nm, em 625/30 nm; ex 640/40 nm, em 690/50 nm; ex 785 nm, em 845/55 nm. Samples were imaged with 1x objective (LaVsion 0.1 NA [WD= 17 mm] or 4x objective (Olympus XLFLUOR 0.28 NA [WD = 10 mm]), and 12x objective (LaVsion 0.53 NA [WD= 10 mm]). Tile scans with 20% or 30% overlap along the longitudinal x-axis and y-axis were obtained using a z-step of 3 μm, 5 μm or 8 μm. Exposure time was 90–120 ms, laser power was adjusted depending on the intensity of the fluorescent signal (in order to never reach the saturation) and the light-sheet width was kept at 80% of maximum.

### Laser-scanning confocal microscopy imaging

After imaging with light-sheet microscopy, areas of interest from the cleared specimens, such as human brain and human kidney, were dissected and imaged with an inverted laser-scanning confocal microscopy (Zeiss, LSM 880) using Zen 2 software (v.10.0.4.910; Carl Zeiss AG). Before imaging, samples were mounted by placing them onto the glass surface of a 35 mm glass-bottom petri dishes (MatTek, P35G-0-14-C) and adding a few drops of BABB to make sure that the imaging region was immersed in BABB [14]. The imaging was done using a 40x oil-immersion objective lens (Zeiss, ECPlan-NeoFluar × 40/1.30 oil DIC M27, 1.3 NA, WD = 0.21 mm). 7.5 × 5 × 1 cm brain section was cut from number 7 labeled and cleared intact human brain slice, and imaged with upright confocal microscopy (MAVIG, RSG4) using Caliber I.D. RS-G4 research software. The working dimensions of stage is 46 × 40 × 66 cm. The images were acquired with UPLFLN 10x objective (OLYMPUS 0.3NA [WD=10mm]). Tile scans were obtained along the longitudinal y-axis. Laser power was adjusted depending on the intensity of the fluorescent signal (in order to never reach the saturation).

### Epifluorescence stereomicroscopy imaging

Cleared 1.5 cm-thick whole human slice was fixed in a glass chamber and was imaged with a Zeiss AxioZoom EMS3/SyCoP3 fluorescence stereomicroscopy using a 1x long working distance air objective lens (Plan Z 1x, 0. 25 NA, WD= 56 mm). The magnification was set as 32x and imaging areas were selected manually to cover half of slice. The images were taken with 405 nm filters and files were exported as tiff images.

### Image processing

Processing, data analysis, 3D rendering and video generation for the imaging data were done on an HP workstation Z840, with 8 core Xeon processor, 196 GB RAM, and Nvidia Quadro k5000 graphics card and HP workstation Z840 dual Xeon 256 GB DDR4 RAM, nVidia Quadro M5000 8GB graphic card. We used Imaris, Arivis, photoshop and Fiji (ImageJ2) for 3D and 2D image visualization. Tile scans were stitched by Fiji’s stitching plugin49. Stitched images were saved in TIFF format and optionally compressed in LZW format to enable fast processing. We removed tiles with acquisition errors using Fiji’s TrakEM2 plugin and Imglib2library. We acquired light-sheet microscopy stacks using ImSpector (v.5.295, LaVision BioTec) as 16-bit grayscale TIFF images for each channel separately. In each pair of neighboring stacks, alignment was done by manually selecting 3 to 4 anatomic landmarks from the overlapping regions, and then the stitching was done sequentially with the Scope Fusion module of the Vision4D (v.2.12.6 × 64, Arivis) software. Landmarks were mainly chosen from the cellular structures on the basis of visual inspection of the anatomical features. After completing the 3D reconstructions, the data visualization was done with Imaris (v.9.3.0, Bitplane) in both volumetric and maximum-intensity projection color mapping. Epifluorescence stereomicroscopy images were saved as series TIFF images. The images were stitched manually in photoshop by overlapping regions from landmarks.

### 3D volume and iso-surface rendering of pig pancreas islets

Insulin positive β-cell volumes were quantified by 3D iso-surfacing using the Imaris software (v.9.3.0, Bitplane, Switzerland). Islet volumes were segmented using the ‘absolute threshold’ thresholding option and the intensity threshold was set manually for most of pancreas with the filter of ‘number of voxels Img=1’. Statistical data parameters (Overall, Area, Center of Homogeneous Mass, Ellipsoid Axis A, Ellipsoid AxisB, Ellipsoid Axis C, Ellipsoid Axis Length A, Ellipsoid Axis Length B, Ellipsoid Axis Length C, Ellipticity (oblate), Ellipticity (prolate), Number of Voxels, Position, Sphericity and Volume) were exported from Imaris to Microsoft Excel (Microsoft, USA) spread sheets (*.xls). Statistical analysis was performed using GraphPad Prism8.

## AUTHOR CONTRIBUTIONS

S.Z. performed the protocol development, organs processing, labeling, clearing, imaging and data analysis. M.I.T. contributed to optimize the protocol, organs processing, imaging and data analysis. H.S. perfused and dissected the human brain and human kidney. E.K. and E.W. generated the INS-EGFP transgenic pig line. E.K. perfused and dissected pig pancreas. J.L. performed the SAXS experiment and data analysis. S.Z. and A.E. wrote the manuscript. All authors commented on the manuscript. A.E. initiated and led all aspects of the project.

## ACKNOWLEDGMENTS

We thank O. Thorn-Seshold for the critical comments on the manuscript, and David Eliezer and Sebastian Doniach for support in the initial stages of this project. This work was supported by the Vascular Dementia Research Foundation, Synergy Excellence Cluster Munich (SyNergy), ERA-Net Neuron (01EW1501A to A.E.), Fritz Thyssen Stiftung (A.E., Ref. 10.17.1.019MN), DFG (A.E., Ref. ER 810/2-1 and TRR127 to E.W. and E.K.), NIH (A.E.), and Helmholtz ICEMED Alliance (A.E.).

## MOVIE LEGENDS

### Supporting Movie 1

#### Beta cells distribution in pig pancreas by SHANEL histology

SHANEL histology revealed individual or groups of beta cells within islets of INS-EGFP transgenic pig pancreas that is several centimeters in size. 3D reconstruction of nanobody boosted EGFP transgenic pancreas imaged by light-sheet microscopy. After SHANEL, all labelled beta cells became evident in the islets throughout the pancreas. In the second part of the movie, segmented EGFP+ islets shown.

### Supporting Movie 2

#### 3D imaging of intact human eye using SHANEL technology

3D reconstruction of intact human eye imaged by light-sheet microscopy. SHANEL technology revealed structural details of sclera, iris and suspensory ligament in intact human eye.

### Supporting Movie 3

#### Abeta plaques accumulation in 92 years large human brain slice by SHANEL histology

Upright confocal microscopy imaging of 0.4 cm thickness of a human brain slice with the cellular resolution (before clearing size: 10 × 7 × 1.5 cm; after clearing size 7.5 × 5 × 1.1 cm). The large scan shows the Abeta plaques are heavily accumulated in specific cortex regions including middle temporal gyrus (MTG) and cingulate gyrus (CG).

### Supporting Movie 4

#### SHANEL histology on centimeters sized human brain tissues for microglia labeling

3D reconstruction of Iba1 antibody labeled microglia cells were evident throughout the centimeters size human brain tissue imaged by light-sheet microscopy.

### Supporting Movie 5

#### SHANEL histology on centimeters sized human brain tissues for neural process labeling

3D reconstruction of Tyrosine hydroxylase (TH) antibody labeled neuronal processes were evident throughout centimeters size human brain tissue imaged by light-sheet microscopy. Individual axonal extension across the gray matter and white matter were visible.

### Supporting Movie 6

#### Cellular and molecular investigation of human kidney using SHANEL histology

3D reconstruction of human kidney cortex showing TRITC-dextran labeled vessels and TO-PRO-3 labeled glomeruli structures, imaged by light-sheet microscopy. Individual glomerulus was tracked by afferent arteriole. Structural profiles of cortex were characterized by the width, glomerular capillary diameter and afferent arteriole diameter.

**Supporting Fig. 1.**
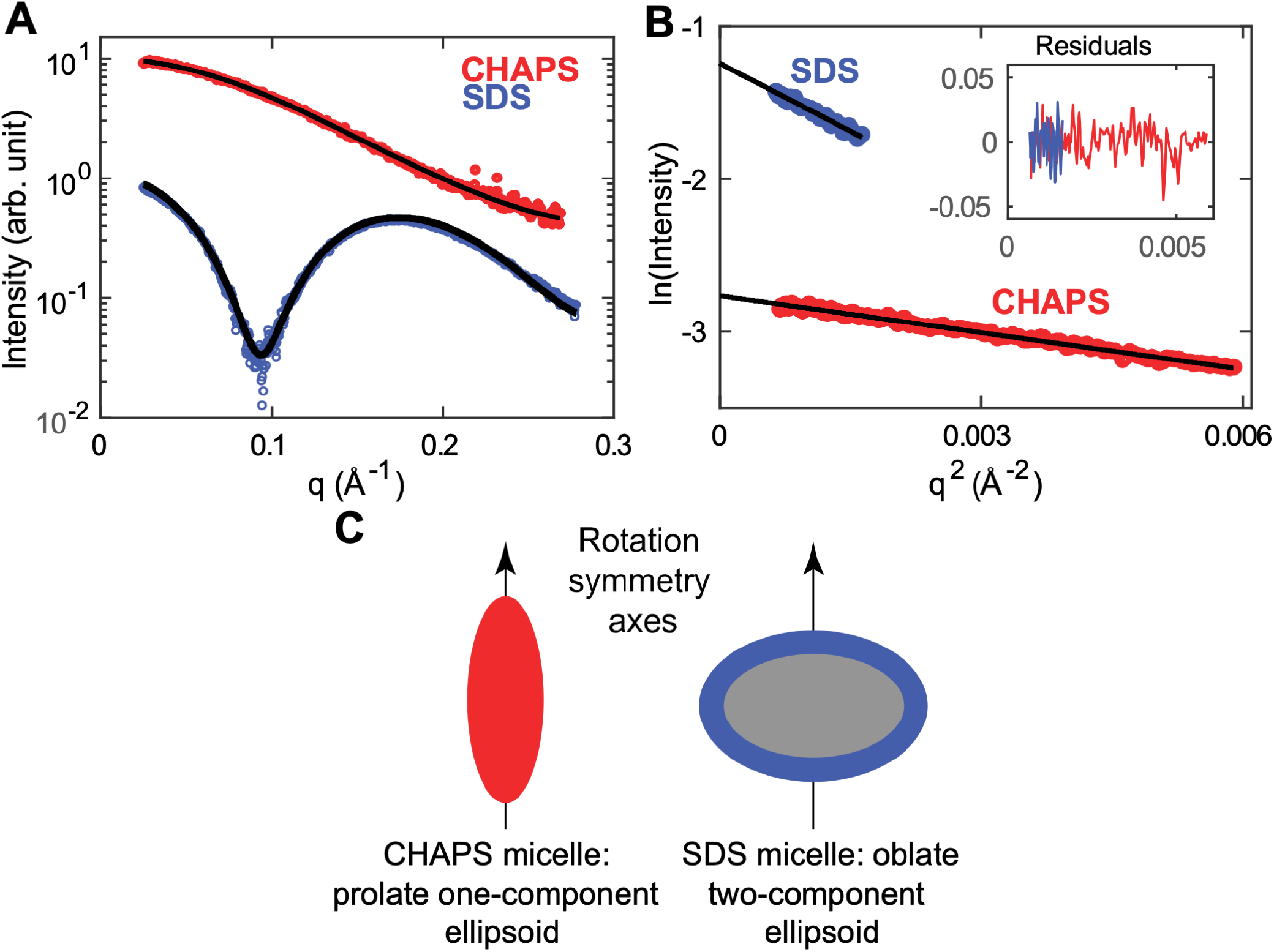
Analysis of detergent micelle shape and size by small-angle X-ray scattering. **(A)** Small-angle X-ray scattering profiles of CHAPS (red) and SDS (blue) micelles in PBS buffer. Fits of ellipsoid to the data are shown in black models (see Methods for details). **(B)** Guinier analysis of the scattering profiles from panel **(A)**. The black lines indicate linear fits to *ln(I)* vs. *q^2^*; the steeper slope for SDS corresponds to a larger radius of gyration *R* The fitting ranges were chosen such that the largest *q*-values are included in the fit, *q_max_*, satify the condition *q* · *R* <.2. The inset shows the residual of the fit, confirming good linearity of the data in the Guinier region. **(C)** Schematics of the geometrical models fit the CHAPS and SDS scattering data, shown to relative scale.

**Supporting Fig. 2.**
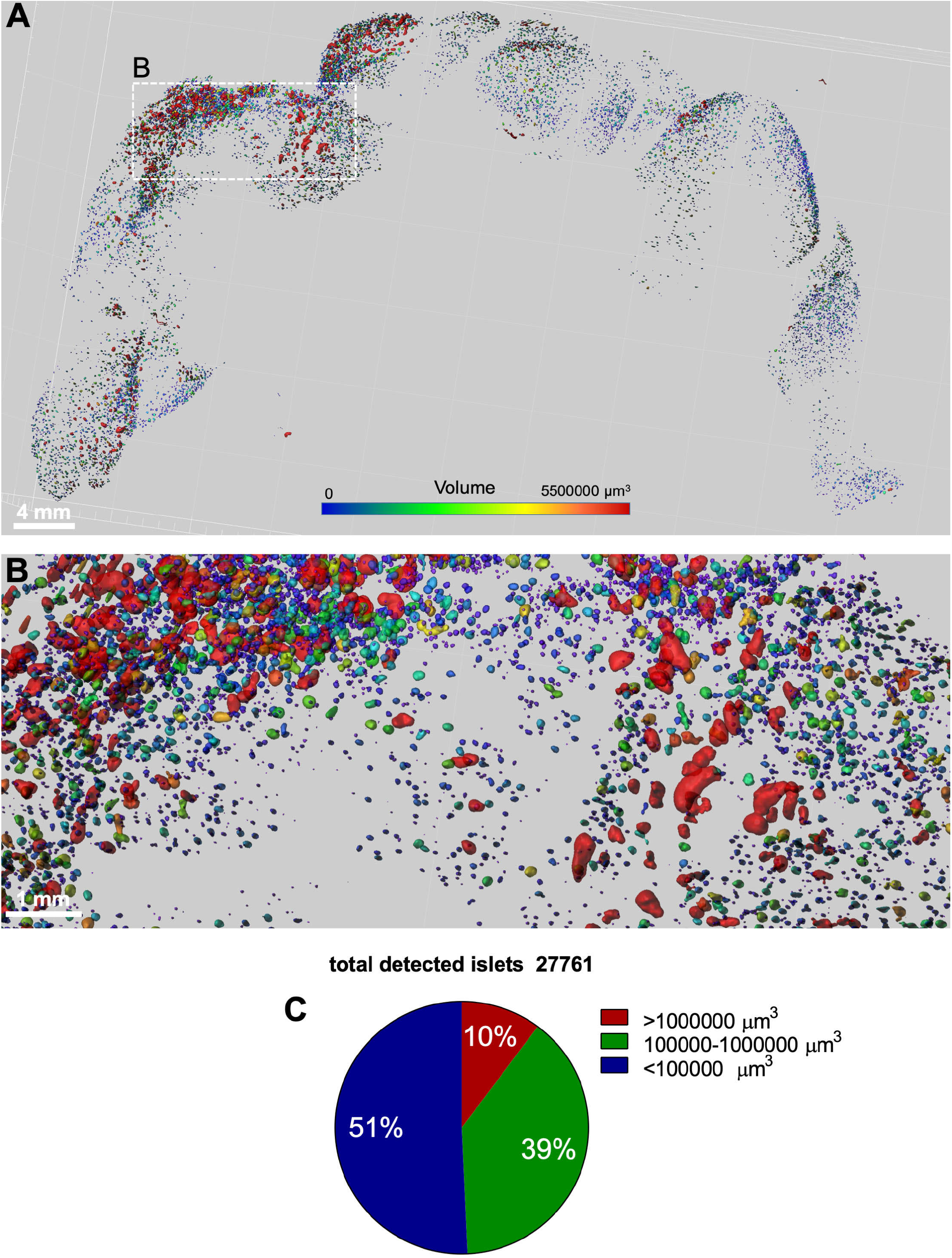
Segmentation and quantification of pancreas islets in transgenic pig after SHANEL histology. *INS*-EGFP transgenic pig pancreas expresses EGFP under the porcine insulin gene *(INS)* promotor labeling beta cells in the islets of Langerhans. We used anti-GFP nanobodies (ref. vDISCO) to enhance and stabilize the signal of EGFP **(A, B)** Demonstration of the 3D distribution of pancreatic beta cells based on their volume in the islets of Langerhans. The heterogeneity of islet sizes is evident. **(C)** Quantification of the total number of detected islets and categorization based on their volume.

**Supporting Fig. 3.**
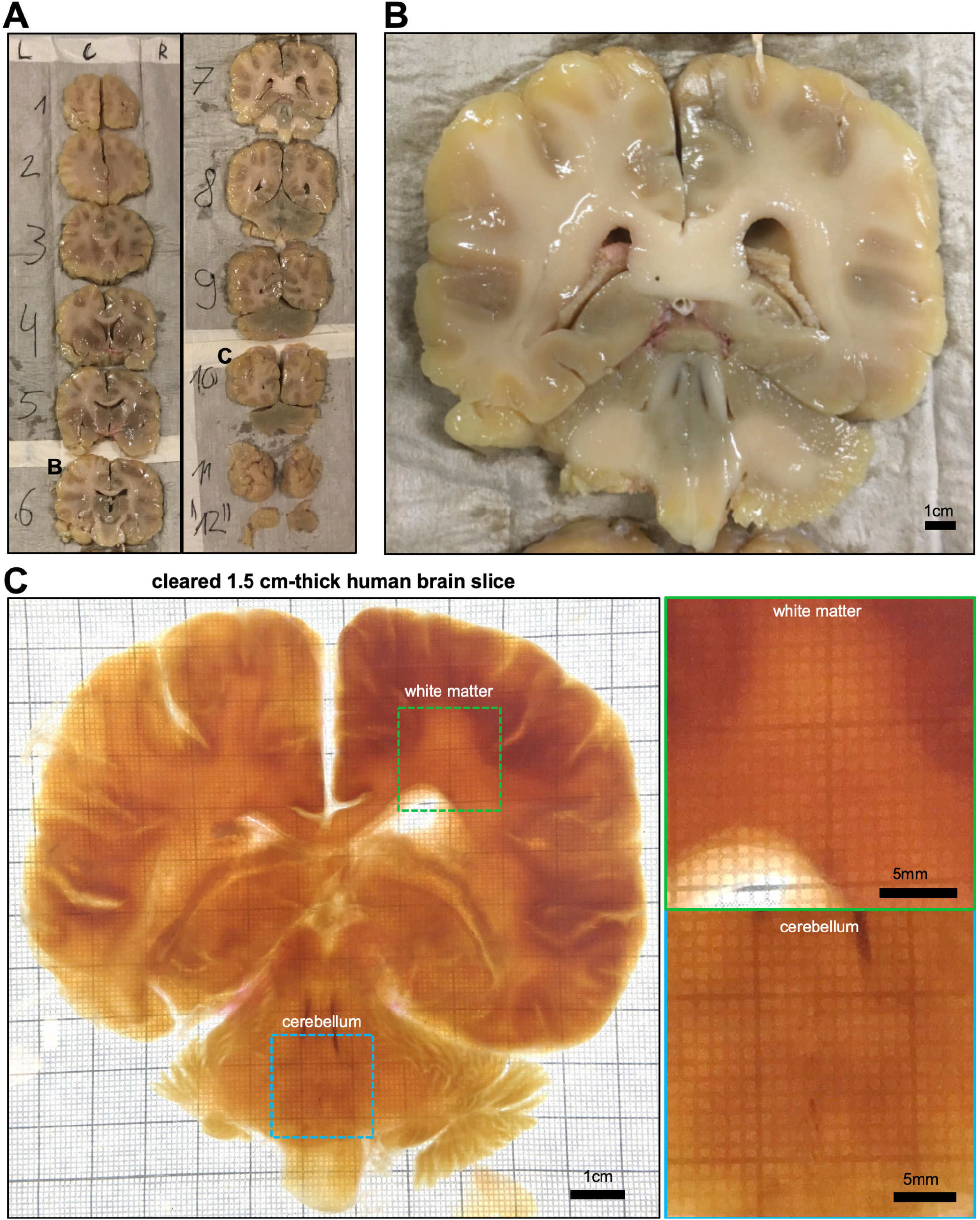
Slicing of intact adult human brain for SHANEL clearing. **(A)** 12 slices of 1.5 cm thickness from an intact adult human brain after CHAPS/NMDEA treatment. **(B)** Photo of an example slice (#7) before SHANEL clearing. **(C)** Photo of the example slice (#7) after SHANEL clearing showing the full transparency of the 1.5 cm thick human brain slice. The colored rectangles are shown in higher magnification on the right hand side showing the heavily myelinated white matter and cerebellum.

**Supporting Fig. 4.**
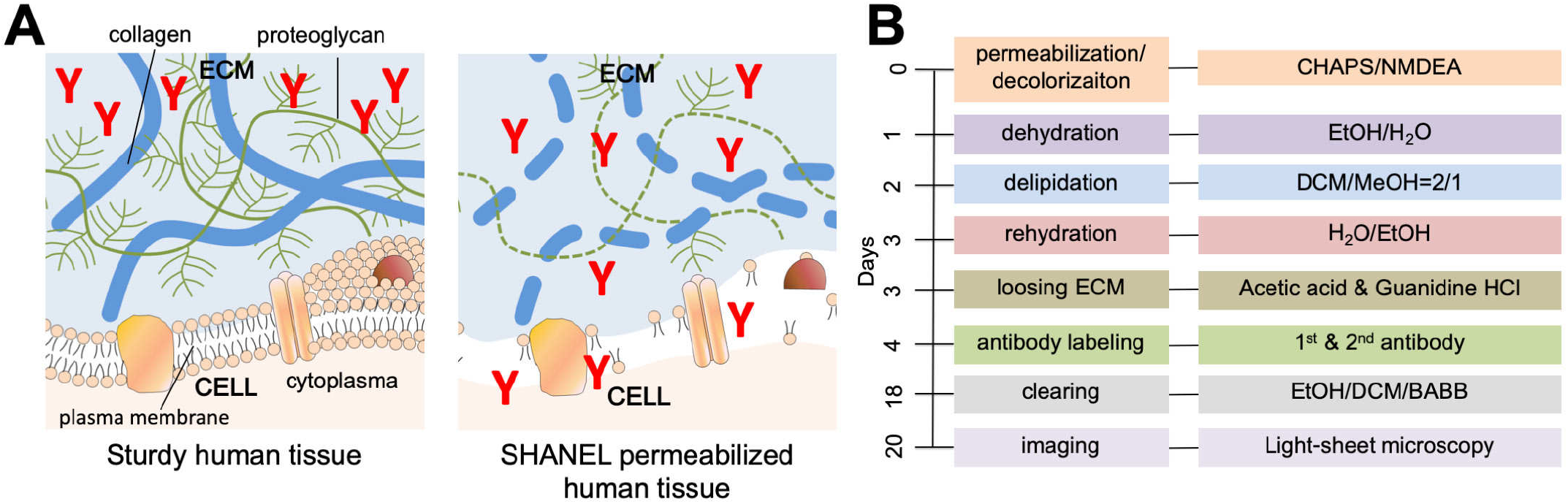
SHANEL histology pipeline. **(A)** SHANEL histology is further characterized by loosening of the extracellular matrix (ECM) using acetic acid & guanidine HCl, which enables antibody-sized molecules to fully penetrate into centimeter-thick sturdy adult human tissues. (B) Step by step SHANEL histology pipeline (with durations) for deep tissue antibody labeling.

**Supporting Fig. 5.**
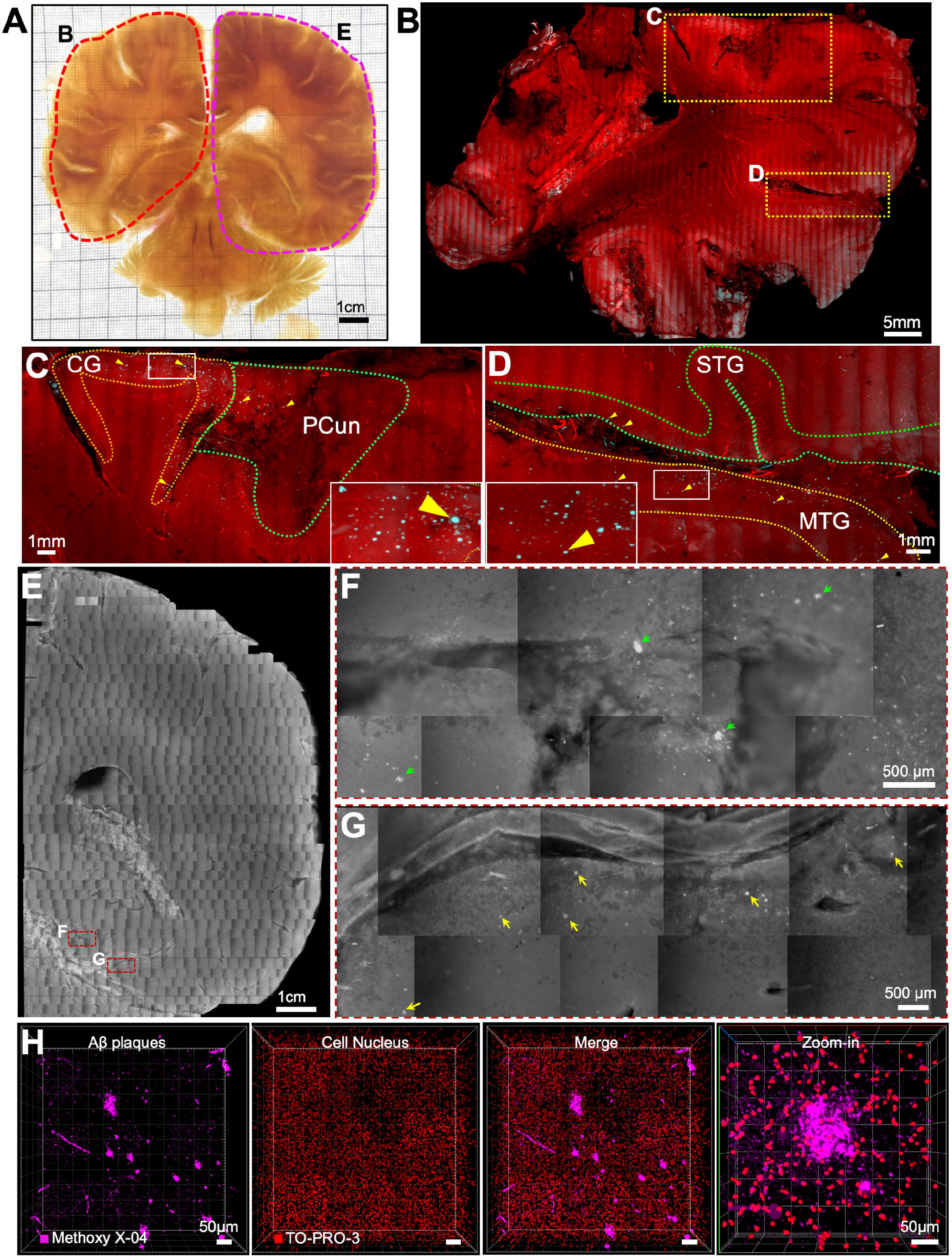
Imaging centimeters sized human brain slice with an upright confocal or with an epifluorescence microscope. **(A)** We imaged large parts of the example slice (#7) using an upright confocal microscope (left side, red region) or an epifluorescence microscope (right side, purple region). **(B-D)** 3D reconstruction of upright confocal images of the left side of the brain in **(A)** showing the TO-PRO-3 labeled cell nuclei (red) and Methoxy-X04 labeled Abeta plaques (cyan). Zoom-in images indicating plaque accumulation regions including cingulate gyrus (CG), precuneus (PCun), superior temporal gyrus (STG) and middle temporal gyrus (MTG) (arrowheads in C and D). **(E-G)** Stitched epifluorescent data of right side of the brain slice in (A) showing Aβ plaques accumulating in the parahippocampal gyrus (PHG, green arrows) and fusiform gyrus (FuG, yellow arrows). **(H)** Tiled 3D confocal images showing Aβ plaques (magenta) and surrounding cell nuclei (red) from the cortex. See also Supporting Movie 3.

**Supporting Fig. 6.**
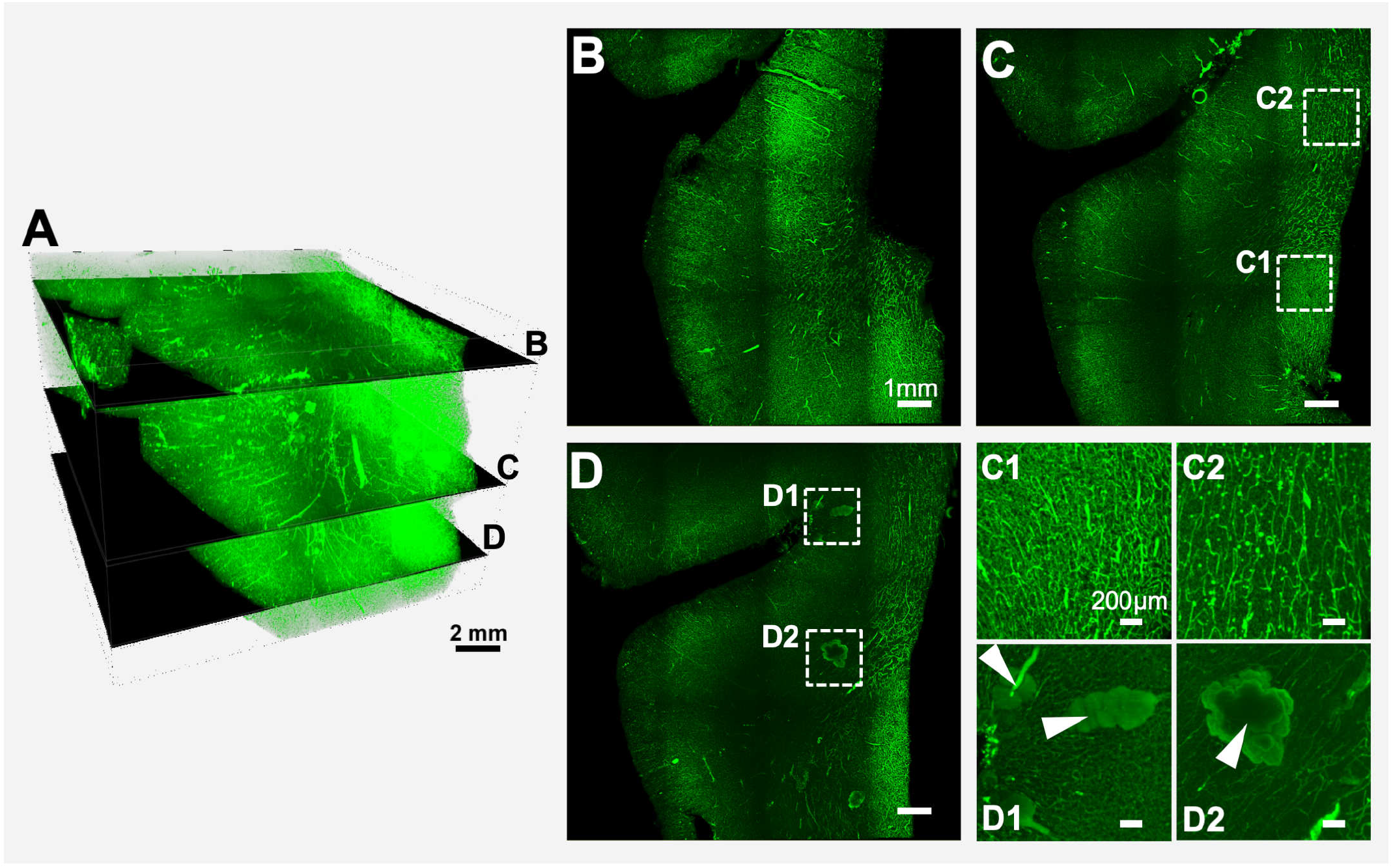
Example of human brain stained with Lectin by SHANEL histology. **(A-D)** Lectin labeling of post-mortem human brain tissue with a size of 3.0 × 1.9 × 1.5 cm. Lectin labeling throughout the entire depth of the centimeters sized human brain tissue is evident. For example, alteration of regional density of vasculature (compare C1 to C2) and tissue abnormalities observed by swollen structures (arrowheads in D1 and D2) are evident.

**Supporting Table 1.** Physical/chemical/geometrical properties of detergent micelles.

| Detergent | CHAPS | SDS | Triton X-100 |
| --- | --- | --- | --- |
| CMC<sup>a</sup> (mM) | 8 Ref. [1] | 1-8 Ref.[1,2] | 0.3 Ref.[3] |
| Aggregation number | 10 Ref. [1]<br>11-12 [from I(0)]<sup>b</sup><br>15-17 [from model]<sup>b</sup> | 80-90 [from I(0)]<sup>b</sup><br>90-100 [from model]<sup>b</sup> | 80 Ref.[4]<br>142 Ref.[5]<br>75-165<sup>c</sup> |
| Monomer mass (Da) | 615 | 288 | 628 Ref.[5] |
| Micelle molecular mass<sup>d</sup> (kDa) | ≈ 7 | ≈ 26 | ≈ 63 |
| Radius of gyration (Å) | 16.0 ± 1.0 Ref.[6]<br>15.5 ± 1.0 | 29.7 ± 1.0 Å | ≈ 33 Å Ref.[4]<br>29.5 ± 0.2 Å Ref.[7] |
| Monomer volume<sup>e</sup> V<sub>mon</sub> (Å<sup>3</sup>) | 830.3 | 414.2 |  |
| Electron density<sup>f</sup> ρ<sub>det</sub> (e/Å<sup>3</sup>) | 0.405 | 0.377 |  |
| Micelle shape | Prolate ellipsoid;<br>Long axis: 30-32 Å<br>Short axis: 10-11 Å | Oblate two-component ellipsoid;<br>Long axis core: 23-25 Å<br>Short axis core: 15-16 Å<br>Outer shell: 2-4 Å |  |

<sup>a</sup>Critical micelle concentration. The CMC for ionic detergents such as SDS depends on ionic strength of the solution[1]. For 150 mM monovalent salt, corresponding to the ionic strength of PBS buffer and the approximate physiological ionic strength the CMC for SDS is 1.5 mM.
<sup>b</sup>Aggregation numbers determined from SAXS analysis. Two independent estimates were obtained by i) analyzing the forward scattering intensity I(0) and ii) fitting the shape of the scattering patterns with a geometrical model (see Methods).
<sup>c</sup>Anatrace catalogue (2014)
<sup>d</sup>Computed from the monomer mass and the estimated aggregation number.
<sup>e</sup>Monomer volumes were computed were calculated from the monomer mass and the published specific densities[2].
<sup>f</sup>Electron densities were computed from the atomic composition and the monomer volume

---

**Supporting Table 2.**
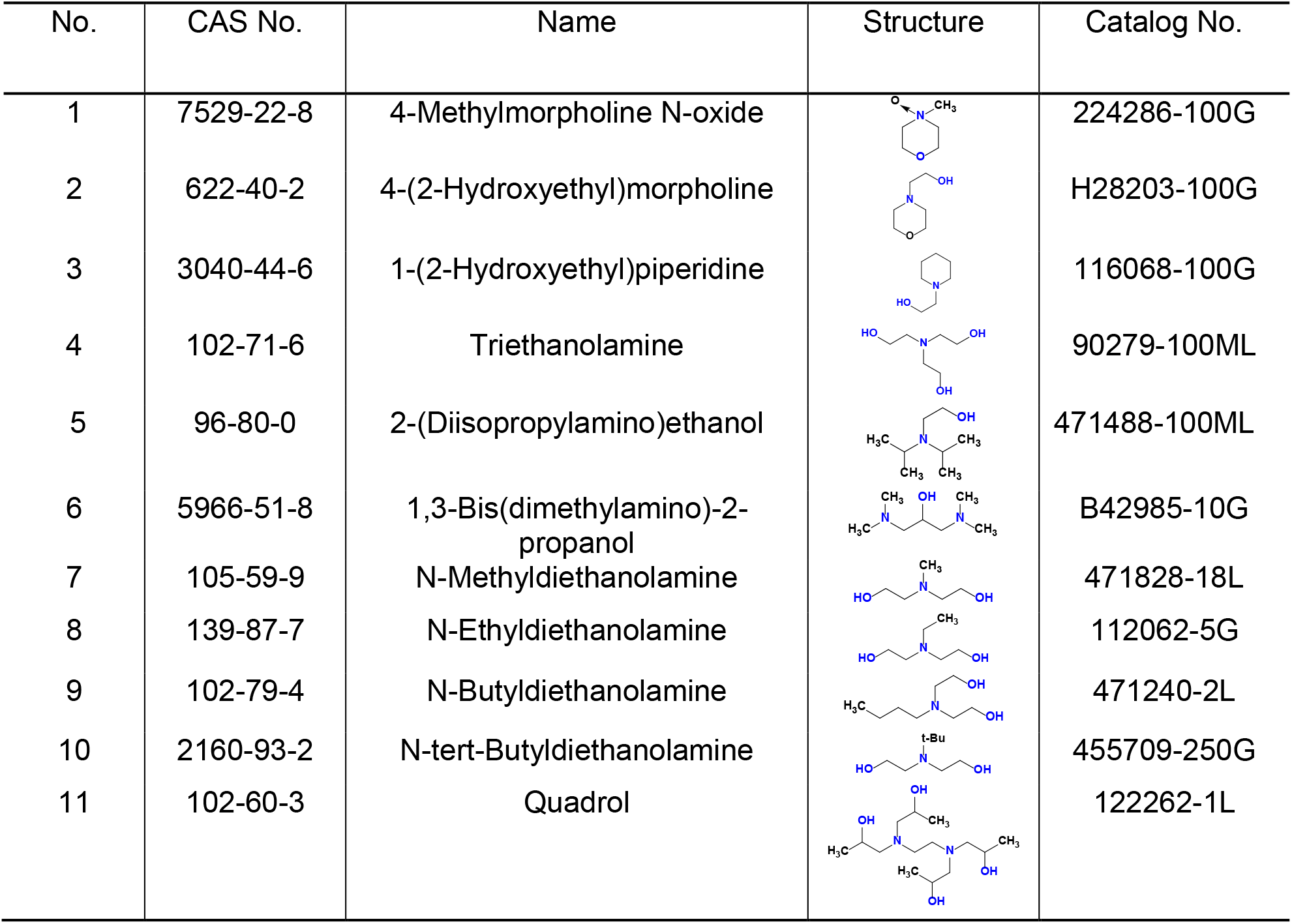
Screening chemicals for blood decolorization.

| No. | CAS No. | Name | Structure | Catalog No. |
| --- | --- | --- | --- | --- |
| 1 | 7529-22-8 | 4-Methylmorpholine N-oxide |  | 224286-100G |
| 2 | 622-40-2 | 4-(2-Hydroxyethyl)morpholine |  | H28203-100G |
| 3 | 3040-44-6 | 1-(2-Hydroxyethyl)piperidine |  | 116068-100G |
| 4 | 102-71-6 | Triethanolamine |  | 90279-100ML |
| 5 | 96-80-0 | 2-(Diisopropylamino)ethanol |  | 471488-100ML |
| 6 | 5966-51-8 | 1,3-Bis(dimethylamino)-2-propanol |  | B42985-10G |
| 7 | 105-59-9 | N-Methyldiethanolamine |  | 471828-18L |
| 8 | 139-87-7 | N-Ethyldiethanolamine |  | 112062-5G |
| 9 | 102-79-4 | N-Butyldiethanolamine |  | 471240-2L |
| 10 | 2160-93-2 | N-tert-Butyldiethanolamine |  | 455709-250G |
| 11 | 102-60-3 | Quadrol |  | 122262-1L |

**Supporting Table 3.** Prices and scalability of screened chemicals.

| Aminoalcohols | 1 | 2 | 3 | 4 | 5 | 6 | 7<sup>a</sup> | 8 | 9 | 10 | 11 |
| --- | --- | --- | --- | --- | --- | --- | --- | --- | --- | --- | --- |
| Package/kg or L | 0.1 | 0.25 | 0.5 | 2.5 | 0.5 | 0.01 | 18 | 0.005 | 2 | 0.25 | 1 |
| €/kg or L | 1670 | 133 | 630 | 72 | 125 | 16300 | 30 | 4740 | 46 | 290 | 71 |
| Color of solution | - | + | + | - | + | + | - | + | - | + | - |

-: colorless; +: color
a: red highlights the chemicals used in SHANEL

**Supporting Table 4.**
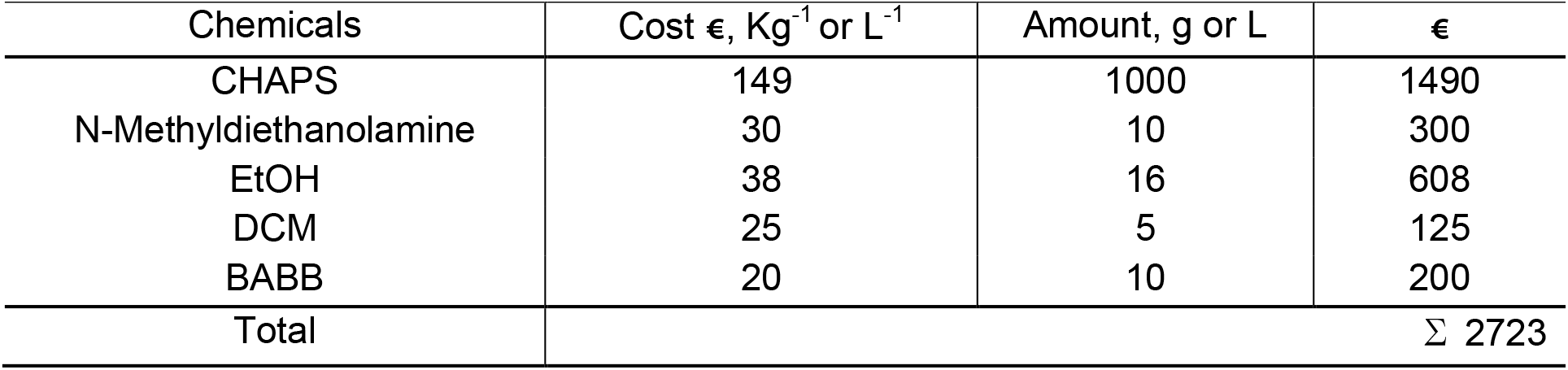
Costs of chemicals for clearing whole adult human brain.

